# SOS1 inhibition enhances the efficacy of and delays resistance to G12C inhibitors in lung adenocarcinoma

**DOI:** 10.1101/2023.12.07.570642

**Authors:** Brianna R Daley, Nancy E Sealover, Erin Sheffels, Jacob M. Hughes, Daniel Gerlach, Marco H Hofmann, Kaja Kostyrko, Barbara Mair, Amanda Linke, Zaria Beckley, Andrew Frank, Clifton Dalgard, Robert L Kortum

**Author notes:** Corresponding Author Robert L. Kortum Uniformed Services University of the Health Sciences 4301 Jones Bridge Rd Bldg C, Rm C2027 Bethesda, MD 20814 **Email:**. **Author Contributions:** BRD, NES, ES, and RLK designed the experiments and analyzed the data; BRD and RLK performed most of the experiments; NES, ES, CD, and DG assisted in performing and analyzing RNA sequencing, JMH assisted with Western blotting, synergy studies, *in vitro* ELDAs, and resistance studies; MHH, KK, and BM assisted in data analysis and interpretation, AL assisted in resistance studies, ZB assisted with Western blotting, BRD and RLK wrote the manuscript, NES, DG, MHH, KK, and BM edited the manuscript. **Competing Interest Statement:** The Kortum laboratory receives funding from Boehringer Ingelheim to study SOS1 as a therapeutic target in *RAS*-mutated cancers.

## Abstract

Clinical effectiveness of KRAS G12C inhibitors (G12Cis) is limited both by intrinsic and acquired resistance, necessitating the development of combination approaches. We found that targeting proximal receptor tyrosine kinase (RTK) signaling using the SOS1 inhibitor (SOS1i) BI-3406 both enhanced the potency of and delayed resistance to G12Ci treatment, but the extent of SOS1i effectiveness was modulated by both SOS2 expression and the specific mutational landscape. SOS1i enhanced the efficacy of G12Ci and limited rebound RTK/ERK signaling to overcome intrinsic/adaptive resistance, but this effect was modulated by SOS2 protein levels. Survival of drug-tolerant persister (DTP) cells within the heterogeneous tumor population and/or acquired mutations that reactivate RTK/RAS signaling can lead to outgrowth of tumor initiating cells (TICs) that drive therapeutic resistance. G12Ci drug tolerant persister cells showed a 2-3-fold enrichment of TICs, suggesting that these could be a sanctuary population of G12Ci resistant cells. SOS1i re-sensitized DTPs to G12Ci and inhibited G12C-induced TIC enrichment. Co-mutation of the tumor suppressor *KEAP1* limits the clinical effectiveness of G12Cis, and *KEAP1* and *STK11* deletion increased TIC frequency and accelerated the development of acquired resistance to G12Ci *in situ*. SOS1i both delayed acquired G12Ci resistance and limited the total number of resistant colonies regardless of *KEAP1* and *STK11* mutational status. These data suggest that SOS1i could be an effective strategy to both enhance G12Ci efficacy and prevent G12Ci resistance regardless of co-mutations.

**Significance:** The SOS1 inhibitor BI-3406 both inhibits intrinsic/adaptive resistance and targets drug tolerant persister cells to limit the development of acquired resistance to clinical KRAS^G12C^ inhibitors in lung adenocarcinoma cells.

## Introduction

Lung cancer is the leading cause of cancer-related death (1). Oncogenic driver mutations in the RTK/RAS pathway occur in 75-90% of lung adenocarcinoma (LUAD), with 30-40% being driven by mutations in *KRAS*. Approximately 13% of LUAD harbor *KRAS^G12C^*mutations (1,2). While KRAS was once considered constitutively active and undruggable, KRAS^G12C^ mutant proteins maintain a rate of hydrolysis sufficient for nucleotide exchange (3–6). This nucleotide cycling between the GTP-bound on and GTP-bound off states, combined with the identification of a unique and targetable binding pocket in KRAS^GDP^(7), afforded the unique opportunity to target KRAS^G12C^ with agents that specifically bind KRAS in the GDP-bound off state and covalently modify the mutated cystine to lock mutant KRAS^G12C^ in the inactive GDP-bound state (7). Pre- clinical development of therapeutics based on this principle showed potent tumor suppression in cellular and animal models (8,9), culminating in the FDA approval of the KRAS^G12C^ inhibitors AMG-510/sotorasib (10,11) and MRTX849/adagrasib (12).

Unfortunately, intrinsic and acquired resistance to both adagrasib and sotorasib limit the clinical benefit in patients with *KRAS^G12C^*-mutated LUAD. Across multiple published clinical trials, previously treated patients receiving adagrasib or sotorasib monotherapy showed response rates of 34-43% (13–15), indicating that intrinsic resistance will limit G12Ci efficacy for more than half of patients with *KRAS^G12C^*-mutated LUAD. Intrinsic G12Ci resistance can be attributed to both pre-existing co-mutations that prevent cancer cells from responding to G12Ci (15–17) and rapid rewiring of intracellular signaling that reactivates RAS effectors to bypass the effects of G12Ci. This reactivation of RAS effector signaling, known as adaptive resistance, is driven both by relief of ERK-dependent negative feedback of RTKs-SOS-RAS signaling (3,18–21) and by pathway activation through newly translated KRAS^G12C^ proteins that are not yet inhibited by G12Ci (22).

Genomic analysis of >400 patients treated with adagrasib or sotorasib found that concurrent loss-of-function (LOF) mutations in *KEAP1*, *SMARCA4*, and *CDKN2A* are associated with inferior clinical outcomes and progression free survival (PFS) ≤ 3 months, indicative of intrinsic resistance (16). Although not associated with inferior overall outcomes, *STK11* mutations were also enriched in patients showing PFS ≤ 3 months (16). *KRAS*-mutated cancers can be broadly categorized into three groups based on co-mutations of the tumor suppressors *TP53*, *CDKN2A*, or *STK11* (23). Of these, tumors with *STK11* mutations show the highest frequency of *KEAP1* mutations (23). Patients with *KRAS* mutant tumors harboring *STK11* co-mutations (24), *KEAP1* co-mutations (25), or both *KEAP1* and *STK11* co-mutations (26) respond poorly to both conventional chemotherapy and immunotherapy. Interestingly, the poor immunotherapy responses for patients whose tumors harbor KEAP1/STK11 co-mutations were only observed for patients with *KRAS*-mutated, but not *KRAS* WT, tumors (26). This finding highlights an important unmet therapeutic need for this patient population.

For patients whose tumors do initially respond to G12Ci, acquired resistance rapidly develops with a mean progression free survival between 5-14 months for patients receiving adagrasib or sotorasib (13–17,27,28). Acquired resistance is often associated with secondary mutations in upstream RTKs, KRAS, NRAS, or downstream RAS effector pathway (4,29,30), although, for roughly half of patients, no definitive driver mutation was observed (4,29,31). As tumors adapt to therapeutic pressure, a proportion of the bulk tumor, known as drug-tolerant persister cells (DTPs) (4,32–34), shows chromatin remodeling that allows cells to survive under drug pressure. These adaptations include broad up-regulation of RTK signaling (35), enhanced ability to handle redox stress (36–40), and entering a near-quiescent state (33–35,41). A subset of the DTP population, known as tumor initiating cells (TICs), has stem-like properties and is capable of asymmetric division and self-renewal. This population likely represents the therapeutic sanctuary of cells responsible for tumor re-growth after the acquisition of additional mutations leading to acquired resistance (4,32,42–45). The idea that the bulk tumor is largely composed of non-TICs is critical to the understanding of acquired resistance, since initial therapeutic efficacy and killing of the bulk population can mask the survival of the small population of TICs that survive and adapt under therapeutic pressure to drive acquired resistance.

SHP2, SOS1, and SOS2 are proximal RTK signaling intermediates critical for activation of both mutated and wild type RAS proteins. The phosphatase SHP2 acts as an adaptor to recruit the RASGEFs SOS1 and SOS2 to the GRB2/GADs complex, allowing for RAS activation at the plasma membrane. While there are currently no SOS2 inhibitors, SHP2i (RMC-4550, SHP099, TNO155) (3,6,18–20,46–49) and SOS1i (BAY-293, BI-3406, and MRTX0902) (50–54) should help overcome intrinsic G12Ci resistance by two distinct mechanisms. First, since G12Cis bind KRAS^G12C^ in the GDP-bound (off) state, SHP2 (3,6,18–20,46–49) and SOS1 (50–54) inhibitors are predicted to enhance G12Ci potency by increasing the amount of KRAS^G12C^ available for G12Ci binding and inhibition. Second, SHP2i (3,6,18–20,46,49), SOS1i (3,6,50,51), or *SOS2^KO^*(55,56) can inhibit adaptive resistance to both G12Ci and MEKi driven by relief of ERK-dependent negative feedback of RTKs-SOS-RAS signaling.

Here, we show that the SOS1i BI-3406 inhibits both intrinsic and acquired G12Ci resistance in LUAD cells. To limit intrinsic resistance, SOS1i both enhanced the efficacy of multiple G12Cis and prevented G12Ci-induced RTK/ERK re-activation associated with adaptive resistance. To limit acquired resistance, SOS1i re-sensitized DTPs to G12Ci and limited survival of G12Ci-induced TICs. SOS1i further delayed acquired G12Ci resistance and limited the total number of resistant colonies, even in cells with *KEAP1* and/or *STK11* inactivation. These data suggest that SOS1i could be an effective strategy to both enhance G12Ci efficacy and prevent G12Ci resistance in *KRAS*-mutated LUAD.

## Results

### SOS1i synergizes with G12Ci to drive transcriptional changes regulating MAPK and hypoxia pathways

G12Cis including ARS-1620, sotorasib, and adagrasib bind KRAS^G12C^ in the GDP-bound state. Thus, secondary treatments that enhance the level of KRAS-GDP should increase the amount of targetable KRAS^G12C^, thereby enhancing the efficacy of G12Ci. SOS1 is a ubiquitously expressed RASGEF and contributes to the continued activation of mutated KRAS^G12C^; SOS1is are currently under investigation in combination with G12Cis in two clinical trials [NCT04975256 and NCT05579092]. We found in *KRAS^G12C^*-mutated H358 LUAD cells heterozygous for the *KRAS^G12C^* allele, the SOS1i BI-3406 synergized with both pre-clinical (ARS-1620) and clinical (sotorasib, adagrasib) G12Cis across a 9 × 9 matrix of SOS1i:G12Ci doses as assessed by an increase in the excess over Bliss across the treatment matrix (**Fig. 1 A-B**). Further deconvoluting the contributions of synergistic potency vs. efficacy showed that the major contribution of SOS1i was to enhance the potency of G12Ci, consistent with the proposed mechanisms of action for both inhibitors (**Fig. 1C**).

**Figure 1.**
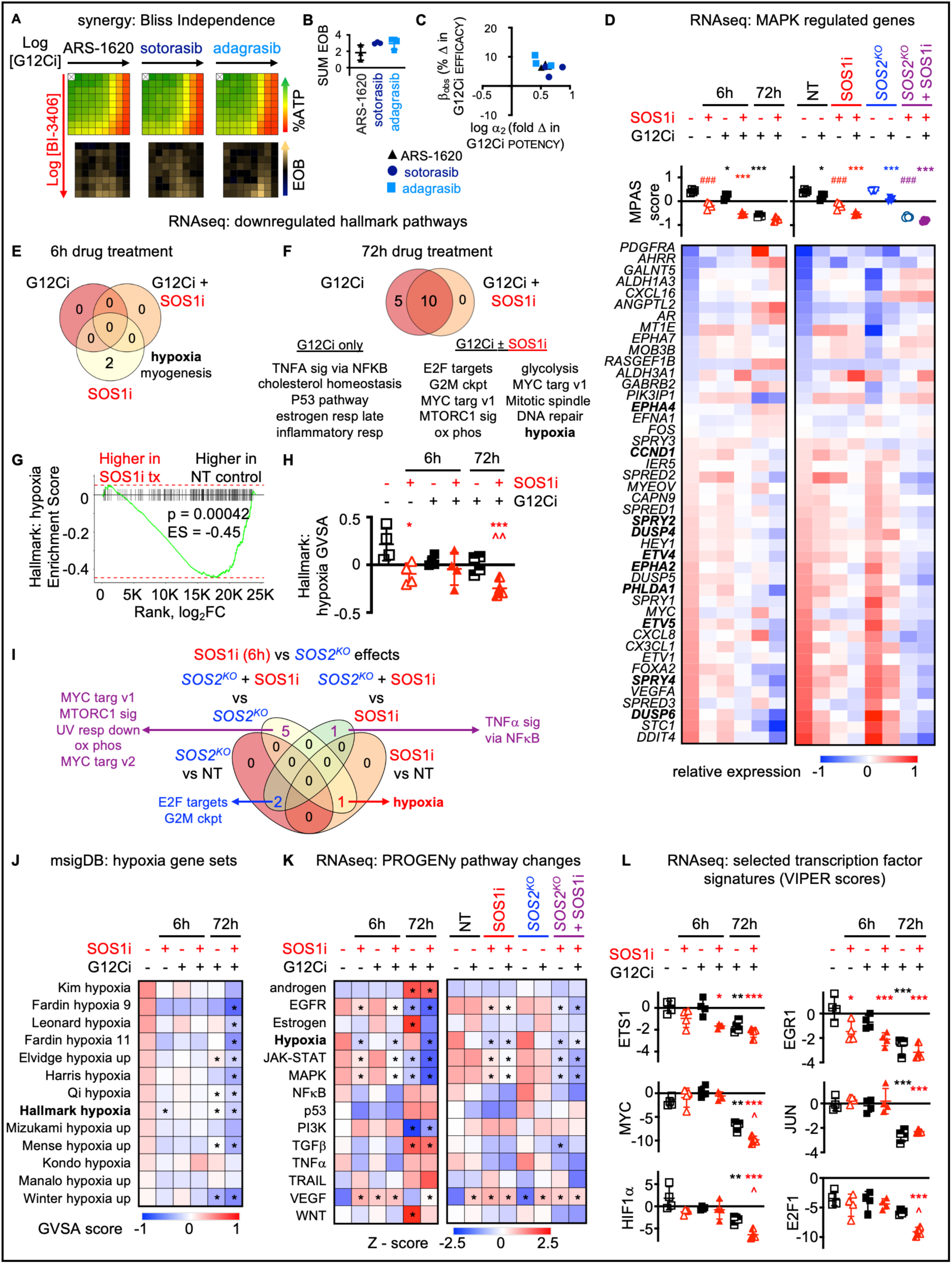
SOS1i synergizes with G12Ci to drive transcriptional changes regulating MAPK and hypoxia pathways. **A.** Heat map of cell viability (top) and excess over Bliss (EOB, bottom) for H358 cells treated with increasing (semi-log) doses of the indicated G12Ci [ARS-1620 (10^-9.5^ – 10^-6^ M), sotorasib (10^-10.5^ – 10^-7^ M), or adagrasib (10^-10.5^ – 10^-7^ M)], the SOS1i BI-3406 (10^-10^ – 10^-6.5^ M) the combination of G12Ci + SOS1i in a 9 × 9 matrix under 3D spheroid culture conditions. Data are the mean from three independent experiments, each experiment had three technical replicates. **B-C**. The sum of excess over Bliss values over the 9 × 9 treatment matrix (**C**) or quantitation of the fold change in G12Ci potency (log α_2_) versus % change in G12Ci efficacy (β_obs_) (**D**) for H358 cells treated with a 9 × 9 matrix of G12Ci + SOS1i from **B**. **D**. MPAS score (top) and heat map of changes in expression of individual MAPK-regulated genes (bottom) from RNA sequencing of 3D spheroid-cultured H358 cells treated with G12Ci (10 nM) ± SOS1i (100 nM) for 6h or 72h (left) or NT or *SOS2^KO^* H358 cells treated with G12Ci (10 nM) ± SOS1i (100 nM) for 6 h (right). Data are averaged from four biologic replicates. * p < 0.05, *** p< 0.001 for G12Ci treated vs. untreated; ### p < 0.001 for SOS1i treated vs. untreated. **E-H**. Venn diagram from GVSA analysis of 50 Hallmark MsigDB gene sets that identified gene sets that were differentially downregulated (p < 0.05) by G12Ci ± SOS1i after 6h (**E**) or 72h (**F**) treatment. GSEA (**G**) and GVSA analysis (**H**) for hypoxia associated genes in H358 cells left untreated (open black square) or treated with a SOS1i for 6h (open red triangle) (**E**), G12Ci ± SOS1i for 6h (closed black square [G12Ci] or red triangle [G12Ci + SOS1i]), or 72h (half-filled black square [G12Ci] or red triangle [G12Ci + SOS1i]) (**F**). **I.** Venn diagram from GVSA analysis of 50 Hallmark MsigDB gene sets that identified nine gene sets that were differentially downregulated (p < 0.05) by SOS1i ± *SOS2^KO^*in H358 cells. **J**. GVSA analysis of MsigDB gene sets that identified hypoxia signatures from RNA sequencing of 3D spheroid-cultured H358 cells treated with G12Ci ± SOS1i for 6h or 72h. Data are averaged from four biological replicates. * p < 0.05 vs. untreated controls. **K**. Heat map of Z-scores for the 14 PROGENy pathway gene sets from RNA sequencing of 3D spheroid-cultured H358 cells treated with G12Ci ± SOS1i for 6h or 72h (left) or NT or *SOS2^KO^* H358 cells treated with G12Ci ± SOS1i for 6 h (right). Data are averaged from four biological replicates. * p < 0.05 vs. untreated controls. **L**. VIPER score for ETS1, EGR1, MYC, JUN, HIF1α, and E2F1 targets from RNA sequencing of 3D spheroid cultured H358 cells untreated (open black square) or treated with a SOS1i for 6h (open red triangle), G12Ci ± SOS1i for 6h (closed black square [G12Ci] or red triangle [G12Ci + SOS1i]) or 72h (half-filled black square [G12Ci] or red triangle [G12Ci + SOS1i]). Data are presented as mean ± from four biologic replicates. * p < 0.05, ** p < 0.01, *** p < 0.001 vs untreated cells; ^ p < 0.05 for G12Ci vs. G12Ci + SOS1i treated for 72h.

We next focused further assessment on the BI-3406:adagrasib combination. We treated H358 cells cultured as 3D spheroids with G12Ci ± SOS1i for 6h or 72h and assessed overall changes in the signaling environment by RNA sequencing. To understand the extent to which additional benefit could be obtained by deletion of the closely related paralog SOS2 (55–58), we further performed RNA sequencing of samples from H358 cells where *SOS2* was deleted using CRISPR/Cas9 compared to non-targeting (NT) controls treated with G12Ci ± SOS1i for 6h. Mutant KRAS signals through the RAF/MEK/ERK cascade to drive proliferation, thus we first assessed the MAPK Pathway Activity Score (MPAS) as a global indicator of on-target transcriptional changes as well as changes in transcription of key MAPK-regulated genes (59). At 6 hours, SOS1i had a more significant impact on MAPK-regulated transcriptional activity compared to G12Ci alone, likely due to the role of SOS1 as a GEF for both mutated and wild-type RAS proteins (**Fig. 1D**). Further, cells treated with the combination of SOS1i + G12Ci showed enhanced inhibition of MAPK-dependent transcriptional changes compared to either drug alone at both 6 and 72h. While *SOS2^KO^* alone did not alter MPAS activity, *SOS2^KO^* decreased MPAS activity in SOS1i-treated samples to a similar extent as G12Ci, confirming a role for SOS2 in regulating MAPK signaling in the absence of SOS1 signaling (**Fig. 1D**).

To globally assess G12Ci ± SOS1i dependent transcriptional changes, we performed gene set enrichment analysis (GSEA) first using the MSigDB hallmark gene set (60) to identify biological processes regulated by SOS1i and G12Ci individually and in combination. At 6h, there were 11 gene sets significantly upregulated and only two gene sets significantly down-regulated by G12Ci and/or SOS1i (**Fig. 1E** and data not shown). At 72h there were no upregulated gene sets and 15 down-regulated gene sets (**Fig. 1F** and data not shown). Further focusing on down-regulated gene sets, we found that pathways associated with hypoxia and myogenesis were significantly downregulated by SOS1i alone (**Fig. 1E**), with hypoxia-associated genes showing a significant enrichment in untreated compared to SOS1i treated samples (**Fig. 1G**). Hypoxia pathways remained downregulated by G12Ci ± SOS1i treatment at 72h (**Fig. 1F**), and gene variant set analysis (GVSA score) showed that hypoxia-regulated genes were more downregulated in SOS1i+G12Ci treated cells compared to cells treated with G12Ci alone at 72h (**Fig. 1H**). Assessment of transcriptional changes in NT vs. *SOS2^KO^*cells either left untreated or treated with SOS1i for 6h identified hypoxia pathways as being regulated by SOS1i but not *SOS2^KO^* (**Fig. 1I**). In contrast, transcriptional changes associated with both E2F targets and the G2M checkpoint were regulated by *SOS2^KO^* but not acute SOS1i treatment, consistent with previous studies showing a role for SOS2 in 3D spheroid growth of *KRAS*-mutated cancer cells (55,56). We further assessed the GVSA for additional MSigDB gene sets associated with hypoxia and found that 9/13 gene sets that were enriched in cells under hypoxic conditions were significantly decreased by SOS1i+G12Ci treatment for 72h, with all 13 showing a more marked decrease in GVSA score for samples treated with SOS1i+G12Ci compared to G12Ci alone (**Fig. 1J**).

We further assessed transcriptional changes in fourteen key cancer-related signaling pathways using PROGENy (Pathway RespOnsive GENes) (61). Transcription of genes associated with EGFR and MAPK signaling was decreased by SOS1i at 6h and either SOS1i or G12Ci at 72h (Fig. 1I). Combined SOS1i + G12Ci further decreased transcription of genes associated with EGFR, JAK-STAT, and MAPK signaling at both 6 and 72h compared to single-drug treated cells, while *SOS2^KO^*showed further pathway inhibition in the presence of a SOS1i (**Fig. 1K**). In contrast, the PROGENy score for genes associated with hypoxia was only significantly decreased in samples from cells treated with a SOS1i (± G12Ci), but not in samples from cells treated with a G12Ci alone (**Fig. 1K**). Assessment of transcription factor signatures associated with early (ETS1, EGR1) or prolonged (MYC, JUN) EGFR/MAPK signaling showed that early transcriptional changes (ETS1, EGR1) were decreased by SOS1i alone at 6h, and this inhibition was enhanced by combined G12Ci + SOS1i at 6 and 72h (**Fig. 1L**). Late EGFR/MAPK regulated transcriptional changes (MYC, JUN) were only downregulated by G12Ci or G12Ci+SOS1i at 72h (**Fig. 1L**). In contrast, gene signatures associated with HIF1α (hypoxia signaling) or E2F1 as a marker of proliferative signaling changes were significantly changed at 72h treatment by combined G12Ci+SOS1i compared to G12Ci alone (**Fig. 1L**). These results suggest a role for SOS1i in regulating both MAPK-dependent signaling associated with G12Ci intrinsic/adaptive resistance and in hypoxia pathways that are associated with the drug-tolerant persister state (62–64).

### SOS1i synergizes with and prevents adaptive resistance to G12Ci

We expanded our evaluation of combined SOS1i + G12Ci in a panel of *KRAS^G12C^*-mutated cell lines. Similar to H358 cells, SOS1i synergistically enhanced the potency of the G12Ci adagrasib in 3D spheroid-cultured H1373 and H23 cells, but not in H1792 or H2030 cells (**Fig. 2A-C and S1A**). Since *SOS2^KO^* increased SOS1i-dependent inhibition of EGFR/MAPK signaling, we diminished the contribution of RTK-SOS2 signaling by treating cells with SOS1i+G12Ci under either low serum conditions (**Fig. S2**) or in *SOS2^KO^* cells (**Fig S3**). In both H1792 and H2030 cells, SOS1i:G12Ci synergy was restored in 2% serum or *SOS2^KO^* (**Fig. 2B-C**). The phosphatase SHP2 is required for full activation of the MAPK pathway and can serve as a proxy for weak combined SOS1/SOS2 inhibition. Similar to combined SOS1i+G12Ci under low serum conditions, SHP2i synergistically enhanced the efficacy of G12Ci in H1792 and H2030 cells (**Fig 2A-C and Fig. S4**). Co-mutations in the tumor suppressors *CDKN2A* and *KEAP1* occur more commonly in patients with *KRAS^G12C^*-mutated tumors showing intrinsic resistance to adagrasib and/or sotorasib (16,29). However, since H1373 and H1792 harbor inactivating *CDKN2A* mutations and H1792, H2030, and H23 harbor inactivating *KEAP1* mutations (Fig. S1B), co-mutational status alone does not result in the variation seen in SOS1i:G12Ci synergy. We therefore assessed the extent to which the relative ratio of SOS1:SOS2 protein abundance determined the requirement for inhibition of SOS2 signaling to promote SOS1i:G12Ci synergy (**Fig. 2D-E**). Indeed, *KRAS^G12C^*-mutated LUAD cell lines showed differential SOS1 vs. SOS2 protein abundance. H1792 and H2030 cells showing higher SOS2 protein abundance compared to H358, H1373, and H23 cells (**Fig. 2D**), and a plot of the relative SOS1:SOS2 ratio versus total EOB for cells treated with SOS1i+G12Ci in 10% serum showed a clear relationship between SOS1:SOS2 protein abundance and SOS1i:G12Ci synergy (**Fig. 2E**). The necessity for combined SOS1/2i to enhance G12Ci efficacy in H1792 and H2030 cells further extended to inhibition of rebound ERK activation following G12Ci treatment. 3D-cultured H358, H1373, H1792, and H2030 cells show rebound ERK phosphorylation 48-72h after G12Ci treatment due to loss of ERK-dependent negative feedback signals (**Fig. 2F-G**). In H358 and H1373 cells that show relatively low SOS2 protein abundance, SOS1i was sufficient to limit rebound ERK activation following G12Ci treatment (**Fig. 2F**). In contrast, in H1792 and H2030 SOS1i alone was not sufficient to block pERK rebound signaling. Instead, combined SOS1i + *SOS2^KO^* was required to limit ERK rebound signaling following three days of G12Ci treatment in these cells. Taken together, these data demonstrate the relative abundance of SOS2 protein determines the extent to which SOS1i can enhance G12Ci efficacy to limit intrinsic/adaptive resistance in *KRAS^G12C^*-mutated LUAD cells.

**Figure 2.**
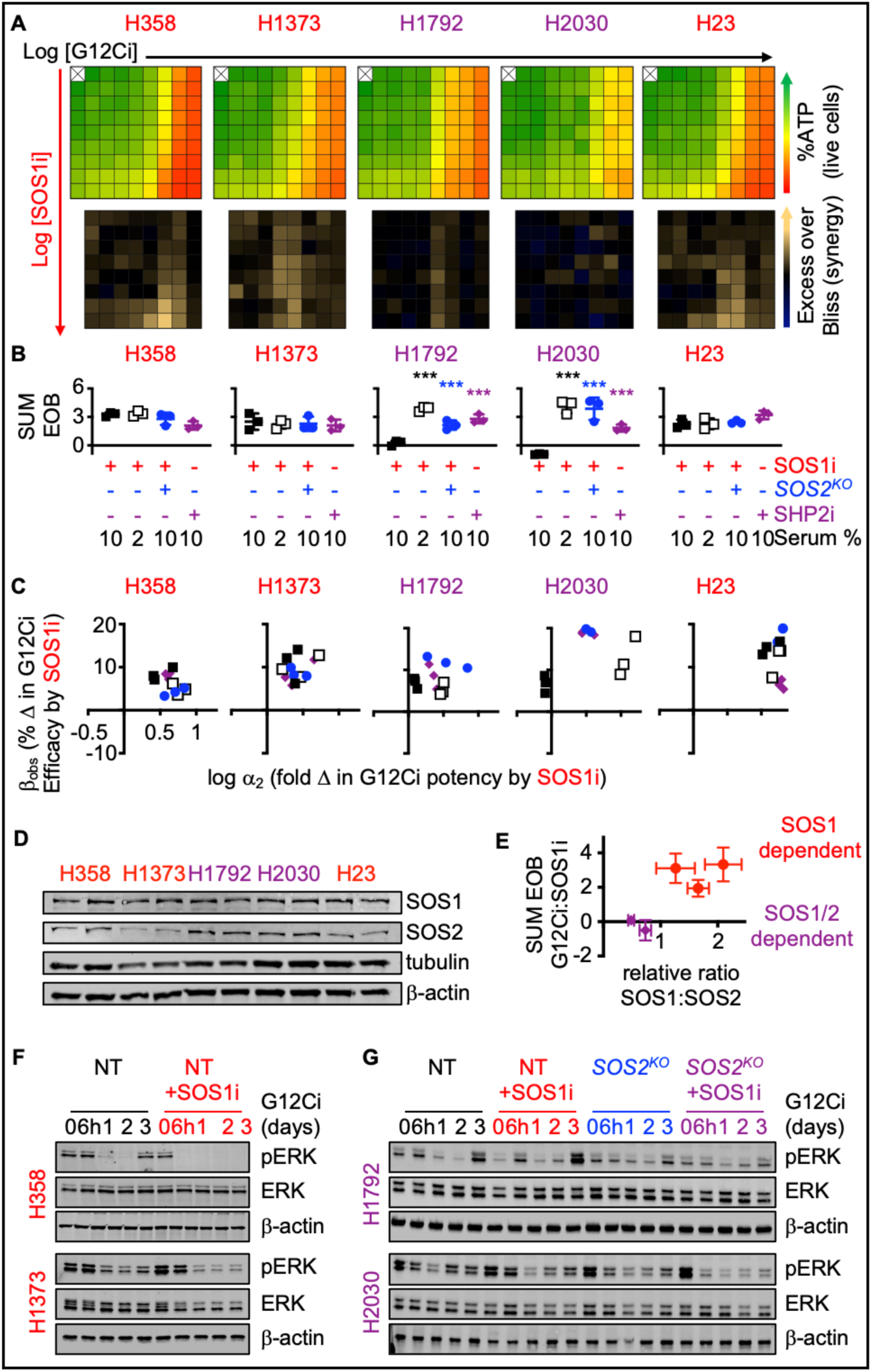
SOS2 expression determines the extent of G12Ci:SOS1i synergy. **A.** Heat map of cell viability (top) and excess over Bliss (EOB, bottom) for the indicated *KRAS^G12C^*-mutated LUAD cell lines treated with increasing (semi-log) doses of the G12Ci adagrasib (10^-10.5^ – 10^-7^), the SOS1i BI-3406 (10^-10^ – 10^-6.5^) or the combination of G12Ci + SOS1i under 3D spheroid culture conditions. Data are the mean from three independent experiments, each experiment had three technical replicates. **B-C.** The sum of excess over Bliss values over the 9 × 9 treatment matrix (**B**) or quantitation of the fold change in G12Ci potency (log αα_2_) versus % change in G12Ci efficacy (β_obs_) (**C**) for the indicated NT (squares) or *SOS2^KO^* (blue circles) cells treated with a 9 × 9 matrix of G12Ci + SOS1i at 10% serum (filled) or 2% serum (open), or treated with a 9 × 9 matrix of G12Ci + SHP2i at 10% serum (purple diamonds). *** p < 0.001 vs. NT cells treated with G12CI + SOS1i in 10% serum. **D-E**. Western blots of whole cell lysates (WCLs) from the indicated LUAD cell lines for SOS1, SOS2, tubulin, and β-actin (**D**) and plot of the relative ratio of SOS1:SOS2 versus the sum of the EOB values for cells treated with a 9 × 9 matrix of G12Ci + SOS1i in 10% serum conditions (**E**). Red circles indicate cell lines showing EOB significantly > 0; purple diamonds indicate cell lines showing EOB ≤ 0. **F-G**. Western blots of WCLs of 3D spheroid-cultured H358 or H1373 cells (**F**) or NT vs. *SOS2^KO^*H1792 or H2030 cells (**G**) treated with G12Ci (10 nM) ± SOS1i (100 nM) for the indicated times. Western blots are for pERK, ERK, and β-actin. Western blots are representative of three independent experiments.

### SOS1i targets G12Ci drug tolerant persister cells

In addition to overcoming intrinsic/adaptive resistance, an optimal therapeutic combination would further delay the development of acquired resistance. Prior to the acquisition of overt secondary mutations, a subset of the bulk tumor can undergo non-genetic adaptations that alter the intracellular redox environment and allow for continued survival in the face of therapeutic pressure (36). This ’drug tolerant persister’ population (DTP) is thought to act as a therapeutic sanctuary for cells to gain additional mutations and develop acquired resistance. Although no single marker fully defines DTP cells, increased aldehyde dehydrogenase (ALDH) activity has been used as a functional marker of cells with increased ability to detoxify the effects of therapy-induced oxidative stress to promote tumor cell survival (37–40). Although DTPs have been well characterized for EGFR-TKIs in LUAD (34,65–70), they have not been well documented after G12Ci treatment. G12Ci treatment resulted in a three-fold increase in the frequency of ALDH^high^ cells in H358 and H1373cells compared to the untreated cells (**Fig. 3A**), suggestive of an enrichment in DTPs.

**Figure 3.**
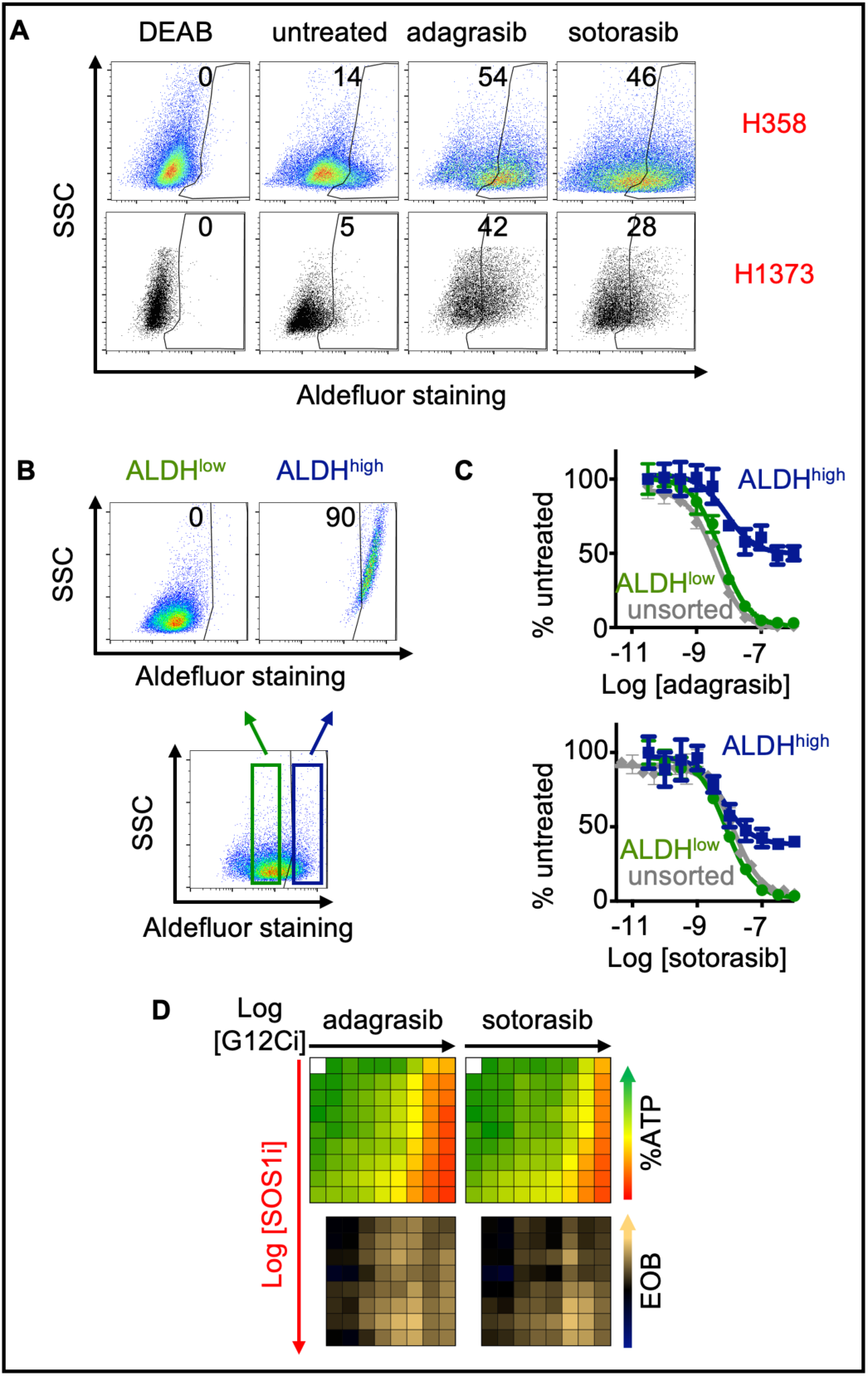
Isolated ALDH^high^ populations are resistant to G12Ci but sensitive to combined G12Ci:SOS1i. **A.** Aldefluor staining for ALDH enzyme activity in the indicated *KRAS^G12C^*-mutated cell lines for DEAB negative control (DEAB), untreated cells, or cells treated with 100 nM adagrasib or sotorasib for 72 hours. **B.** Aldefluor staining for ALDH enzyme activity in H358 cells and gaiting strategy for isolating ALDH^low^ (green) versus ALDH^high^ (dark blue) populations. **C.** G12Ci dose response curves for unsorted (grey), ALDH^low^ (green), and ALDH^high^ (dark blue) H358 cells treated with increasing doses of adagrasib (left) or sotorasib (right). Data are the mean ± sd from three independent experiments, each experiment had three technical replicates. **D.** Heat map of cell viability (top) and excess over Bliss (EOB, bottom) for the ALDH^high^ H358 cells treated with increasing (semi-log) doses of the G12Ci adagrasib (left) or sotorasib (right) (10^-11^ – 10^-7.5^), the SOS1i BI-3406 (10^-10^ – 10^-6.5^) or the combination of G12Ci + SOS1i under 3D spheroid culture conditions in ALDH^high^ sorted H358 cells. Data are the mean from three independent experiments, each experiment had three technical replicates.

Because therapeutic resistance is thought to arise out of the DTP population, we further assessed the extent to which SOS1i could target G12Ci ALDH^+^ DTPs. H358 cells were sorted into ALDH^high^ versus ALDH^low^ populations and assessed for G12Ci sensitivity compared to unsorted populations (**Fig. 3B**). While unsorted and ALDH^low^ cells demonstrated similar sensitivity to increasing doses of G12Ci treatment, adagrasib and sotorasib potency were markedly reduced in sorted ALDH^high^ populations (**Fig. 3C**), showing >30% survival. However, despite this relative insensitivity to G12Ci alone, SOS1i + G12Ci showed marked synergy in sorted ALDH^high^ cells (**Fig. 3D**). These data suggest that SOS1i can re-sensitize DTPs to G12Ci and potentially target pre-resistant LUAD populations prior to the development of acquired resistance.

Tumor initiating cells (TICs) are a subset of the drug tolerant persister population with stem-like properties that are capable of self-renewal and asymmetric division; the TIC population is thought to represent the sanctuary population responsible for tumor recurrence after treatment failure (42,44). HIF1α promotes transcription of genes responsible for TIC activity (71,72) and hypoxia signatures are associated with cancer stemness (62–64) and poor survival for patients with LUAD (73,74). As SOS1i ± G12Ci inhibited hypoxia gene signatures and HIF1α dependent transcription (**Fig. 1**), we hypothesized that SOS1 may regulate TIC survival. TIC frequency can be modeled *in situ* by assessing the frequency with which cells can grow as 3D spheroids from a single cell (tumor initiating cells; TICs). G12Ci treatment (72h) caused a 2-3-fold increase in TIC frequency in H358, H1373, H1792, and H2030 cells (**Fig. 4A**), indicating that tumor-promoting DTPs are enriched in G12Ci treated populations. Since SOS1i (± *SOS2^KO^*) inhibited G12Ci adaptive resistance (**Fig. 2**), we assessed the ability of SOS1i to target G12Ci-induced TICs. In both H358 and H1373 cells, TIC frequency was inhibited by SOS1i in a dose-dependent manner, illustrating the dependency of TIC outgrowth on SOS1 signaling (**Fig. 4B and S5**). Further, TICs were enriched in sorted ALDH^high^ populations, and survival of ALDH^high^ TICs was inhibited by SOS1i (**Fig. 4C**). Unlike H358 and H1373 cells, TIC frequency in H1792 and H2030 cells was not decreased by SOS1i alone, but required combined SOS1i + *SOS2^KO^* indicating the importance of SOS2 to TIC survival in these cell lines (**Fig. 4D**).

**Figure 4.**
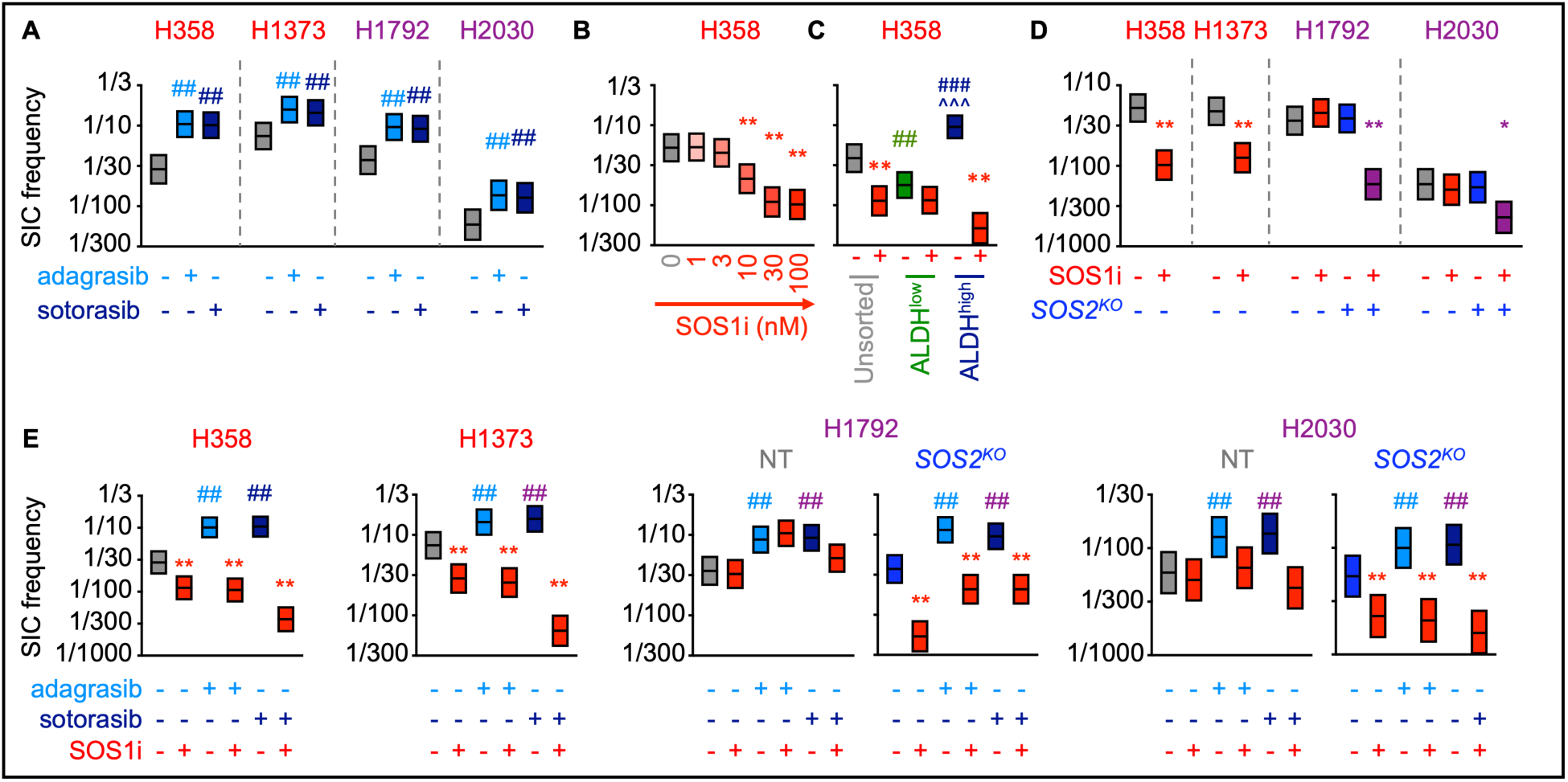
SOS1i ± *SOS2^KO^* prevents G12Ci-induced TIC outgrowth. **A.** TIC frequency from *in situ* extreme limiting dilution assays (ELDAs) of the indicated cell lines pre-treated with 100 nM adagrasib or sotorasib for 72 hours. ## χ^2^ < 0.01 vs. untreated for SIC upregulation. **B.** TIC frequency from *in situ* ELDAs of H358 cells treated with the indicated SOS1i doses. **\*** χ^2^ < 0.05, ** χ^2^ < 0.01 vs. NT untreated. **C.** TIC frequency from *in situ* ELDAs in unsorted (grey), ALDH^low^ (green), and ALDH^high^ (dark blue) H358 cells left untreated or treated with 100 nM BI-3406 (SOS1i). **χ^2^ < 0.01 vs untreated; ## χ^2^ < 0.01, ### χ^2^ < 0.001 vs. unsorted cells; ^^^ χ^2^ < 0.001 vs. ALDH^low^ cells. **D.** TIC frequency from *in situ* ELDAs in the indicated NT or *SOS2^KO^* LUAD cell lines treated with 100 nM BI-3406 (SOS1i). * χ^2^ < 0.05, ** χ^2^ < 0.01 vs. NT untreated. **E.** TIC frequency from *in situ* ELDAs of the indicated NT or *SOS2^KO^* cell lines pre-treated with adagrasib or sotorasib for 72 hours to upregulate TICs, and then left untreated or treated with BI-3406. Data are representative of three independent experiments.

Since SOS1i (± *SOS2^KO^*) inhibited G12Ci adaptive resistance (**Fig. 2**), we assessed the ability of SOS1i to target G12Ci-induced TICs. In both H358 and H1373 cells, SOS1i inhibited G12Ci-induced TIC outgrowth (**Fig. 4E**). In contrast, while H1792 and H2030 cells showed a similar 3-fold increase in TIC frequency post G12Ci treatment (Fig. 4A), SOS1i + *SOS2^KO^*was necessary to effectively inhibit survival of G12Ci-induced TICs (Fig. 4E). These data illustrate that inhibition of proximal RTK signaling can target G12Ci-induced DTPs and TIC survival, thereby potentially reducing the pool of cells capable of promoting acquired resistance.

### SOS1 inhibition limits G12C inhibitor resistance in *KRAS^G12C^*-mutated cells

To directly assess the ability of SOS1i to limit the development of G12Ci acquired resistance, we performed multi-well *in situ* resistance assays (75) in *KRAS^G12C^*-mutated LUAD cells treated with G12Ci ±SOS1i over 6-12 weeks. Here, cells are plated at low density in the inner 60 wells of a 96-well plate and treated with a single dose (or combination) of drug(s). Wells are re-fed weekly and wells showing >50% confluence are scored as resistant to that dose of drug(s). We previously showed that while low doses of RTK/RAS pathway targeted inhibitors cause a proliferative delay, acquired resistance to RTK/RAS pathway inhibitors including G12Cis can be modeled using a ≥ EC_80_ dose of G12Ci (76). For H358 cells, 10 nM adagrasib modeled G12Ci acquired resistance whereas 1, 3, or 6 nM caused a slight delay in proliferation but still showed intrinsic resistance. To determine both whether SOS1i could reduce the G12Ci dose needed to overcome intrinsic resistance and the extent to which SOS1i could delay or reduce the development of G12Ci acquired resistance, we treated H358 cells with 1, 3, 6, or 10 nM adagrasib ± increasing doses of SOS1i (10 – 300 nM) (**Fig. S6A**). While SOS1i had minimal effects at 1 nM G12Ci, SOS1i caused a dose-dependent delay in G12Ci outgrowth and a reduction in the overall fraction of H358 cell cultures showing G12Ci resistance over 12 weeks of treatment. Indeed, at an intermediate (6 nM) adagrasib dose that only delayed H358 outgrowth by 1-2 weeks, SOS1i (100nM) inhibited the development of acquired G12Ci resistance in 80% of cultures, and completely inhibited the development of resistance to 10 nM adagrasib (**Fig. S6A**). We then modeled adagrasib and sotorasib acquired resistance in H358, H1373, H1792, and H2030 cells at increasing doses of G12Ci ± SOS1 (**Fig. S6B-C**) and at the maximum G12Ci dose with SOS1i ± *SOS2^KO^* (**Fig. 5A-B**). In all four cell lines, SOS1i reduced the frequency of cultures showing G12Ci resistance (**Fig. S6**), even at G12Ci doses sub-optimal for modeling G12Ci resistance (outgrowth at < 6 weeks). For both H1792 and H2030 cells, G12Ci doses > 1 μM were required to delay outgrowth, and we were unable to model sotorasib resistance in H2030 cells (**Fig. 5B**). At maximal G12Ci doses for modeling acquired resistance, 100 nM SOS1i completely inhibited acquired G12Ci resistance in H358 and H1373 cells either by delaying the time to develop resistance and/or by reducing the overall percentage of G12Ci resistant cultures in H1792 and H2030 cells (**Fig. 5**). Further, in H1792 and H2030 cells, the ability of SOS1i to limit G12Ci acquired resistance was enhanced by using either a higher dose SOS1i (300 nM) or by combining with *SOS2^KO^* (**Fig. 5**). These data suggest that SOS1i both prolongs the window of G12Ci efficacy and limits the development of G12Ci acquired resistance.

**Figure 5.**
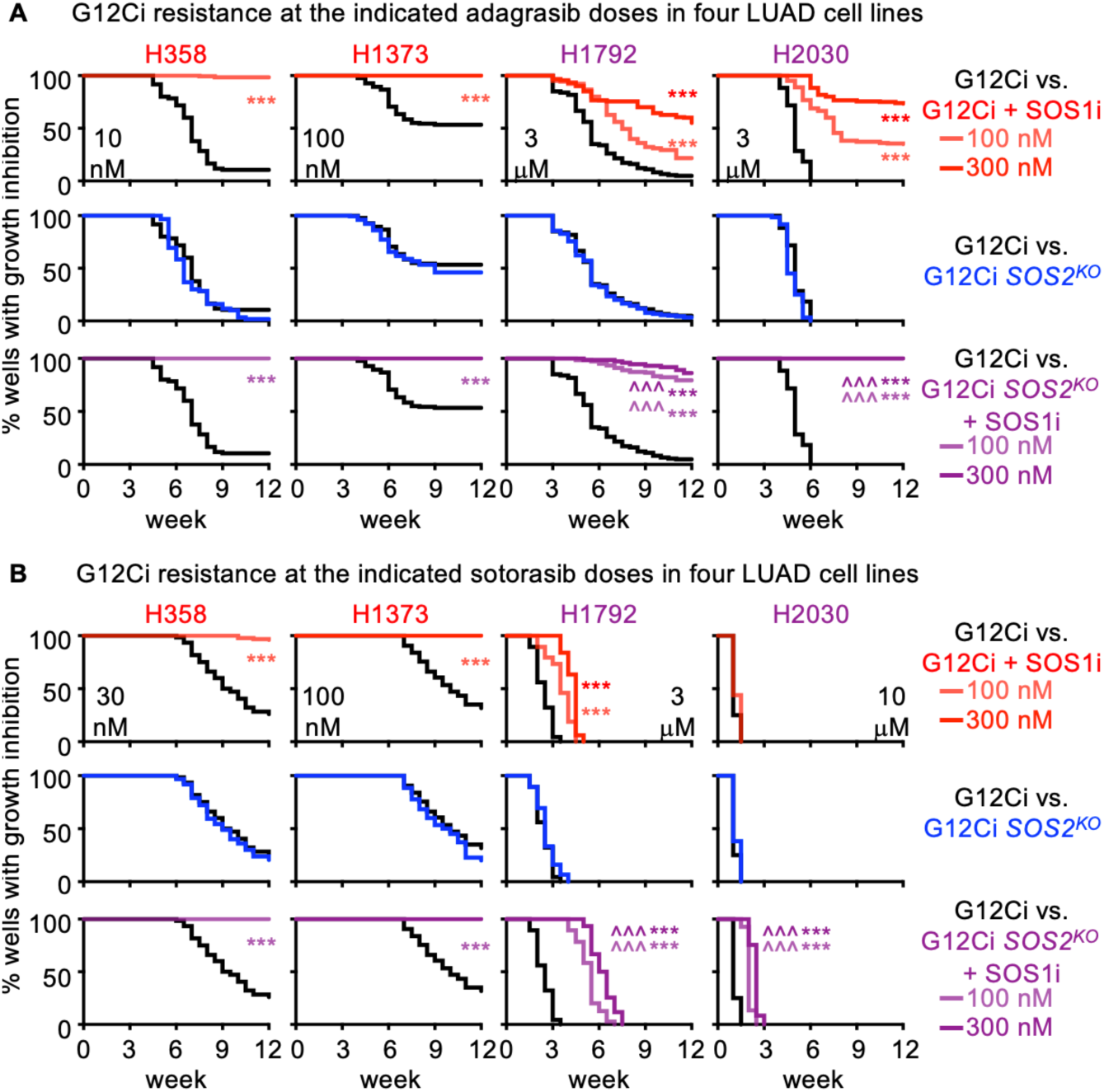
SOS1 inhibition limits the development of acquired G12Ci resistance. Multi-well resistance assay were performed as outlined in the Materials and Methods. **A-B.** G12Ci resistance to the indicated dose of adagrasib (**A**) or sotorasib (**B**) NT or *SOS2^KO^* cells H358, H1373, H1792, or H2030 cells treated with G12Ci alone (NT black; *SOS2^KO^* blue) or G12Ci + SOS1i (NT red; *SOS2^KO^*purple) at 100 nM (lite) or 300 nM (dark). Data are pooled from three independent experiments. *** p < 0.001 vs. G12Ci alone; ### p < 0.001 for cells treated with 100 vs 300 nM SOS1i; ^^^ p < 0.001 for NT vs. *SOS2^KO^* cells treated with SOS1i.

### SOS1i targets DTPs in *KEAP1^KO^* cells to limit acquired G12Ci resistance

H1792 and H2030 cells harbor inactivating co-mutations in the tumor suppressors *KEAP1* and/or *STK11* (**Fig. S1B**) and these co-mutations have been associated with a poorer response to G12Ci treatment and worse overall prognosis for patients (16,29,77,78). However, whether the differential responsiveness of *KRAS^G12C^*-mutated cell lines to G12Ci + SOS1i is directly related to inactivation of *KEAP1* or *STK11* is difficult to assess due to the genetic diversity between different cell lines and the relatively low numbers of KRASG12C mutated LUAD cell lines available for study. To directly assess the impact of *KEAP1* or *STK11* co-mutations to G12Ci + SOS1i combination therapy, we used a panel of isogenic H358 cells where *KEAP1* and *STK11* had been deleted individually or in combination (79). We first assessed the importance of these co-mutations in altering G12Ci:SOS1i synergy after short-term (72h) treatment of 3D spheroids; while *KEAP1^KO^*did not alter G12Ci:SOS1i synergy, *STK11^KO^* showed an apparent decrease in drug-drug synergy, although the overall decrease in cell number at high G12Ci + SOS1i doses did not appear appreciably different (**Fig. 6A**). Intriguingly, *STK11^KO^* cells were more sensitive to G12Ci alone, causing a ½ log decrease in the EC_50_ for adagrasib (**Fig. 6B-C**) which led to an apparent decrease in the excess over bliss value (**Fig. 6D**).

**Figure 6.**
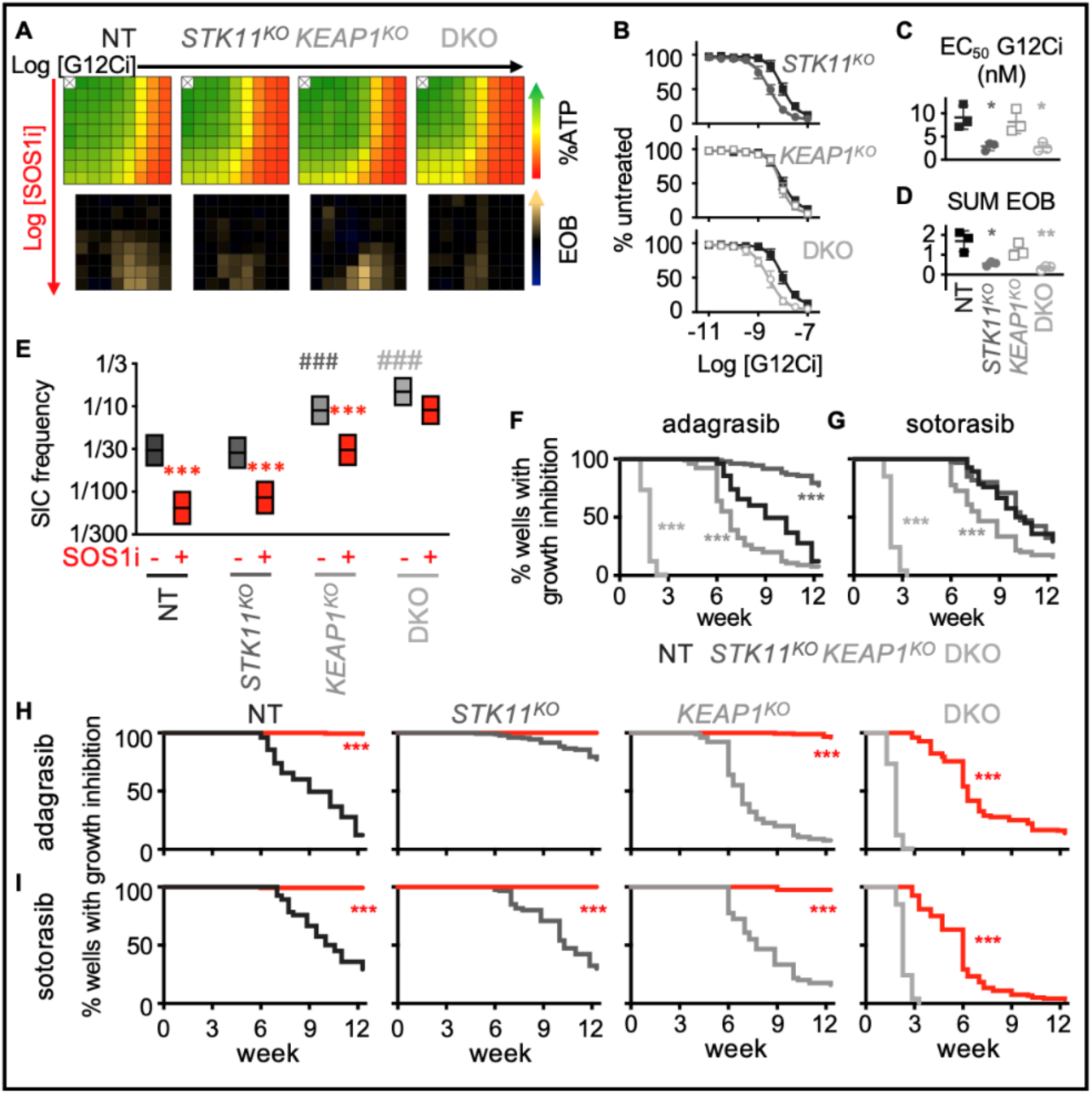
*KEAP1* and *STK11* co-mutations regulate resistance to G12Ci + SOS1i. **A-D.** Heat map of cell viability (top) and excess over Bliss (EOB, bottom) for H358 NT cells or H358 cells where *STK11* and/or *KEAP1* were deleted treated with increasing (semi-log) doses of the G12Ci adagrasib (10^-11^ – 10^-7.5^), the SOS1i BI-3406 (10^-10^ – 10^-6.5^) or the combination of G12Ci + SOS1i for four days under 3D spheroid culture conditions (**A**), adagrasib dose response curves (**B**) EC_50_ values (**C**) in NT (black closed squares), *STK11^KO^* (dark grey closed circles), *KEAP1^KO^* (grey open squares), and *STK11/KEAP1 DKO* (light grey open circles) cells in the absence of SOS1i, and sum of excess over Bliss values over the 9 × 9 treatment matrix from A (**D**). * p < 0.05, p < 0.01 vs. NT cells treated with G12Ci + SOS1i. Data are the mean from three independent experiments, each experiment had three technical replicates. **E**. TIC frequency from *in situ* ELDAs in the indicated H358 cell lines left untreated or treated with 100 nM BI-3406 (SOS1i). ** χ^2^ > 0.01, *** χ^2^ < 0.001 for SOS1i treated cells vs. untreated controls for each cell line; ### χ^2^ < 0.001 s vs. NT untreated cells; ^^ χ^2^ < 0.01 for *STK11^KO^* vs. either *KEAP1^KO^* or *STK11/KEAP1* DKO cells. **F-G**. Multi-well *in situ* resistance assays comparing the development of G12Ci adagrasib (**F**) or sotorasib (**G**) resistance between NT (black), *STK11^KO^* (dark grey), *KEAP1^KO^* (grey), and *STK11/KEAP1 DKO* (light grey) H358 cells. *** p < 0.001 compared to NT cells. **H-I**. Multi-well *in situ* resistance assays comparing the development of G12Ci adagrasib (**H**) or sotorasib (**I**) resistance in NT (black), *STK11^KO^* (dark grey), *KEAP1^KO^* (grey), and *STK11/KEAP1 DKO* (light grey) cells treated with G12Ci alone () or in the presence of SOS1i BI-3406 (300 nM, red lines). *** p < 0.001 for G12Ci + SOS1i vs. G12Ci treated cells.

To determine whether *KEAP1* and/or *STK11* co-mutations affected the survival of DTPs and development of acquired G12Ci resistance, we assessed TIC frequency. Both *KEAP1^KO^* and *KEAP1/STK11 double knock-out (DKO)* cells showed an increased TIC frequency compared to NT and *STK11^KO^* cells (**Fig. 6E**). The increased TICs in *KEAP1^KO^* were significantly inhibited by SOS1i, but not to levels seen in either NT or *STK11^KO^* cells suggesting that KEAP1 mutations alter resistance to combined G12Ci + SOS1i. To directly test this possibility, we performed long-term *in situ* resistance assays to both adagrasib and sotorasib in the panel of H358 KO cells.

When assessing G12Ci resistance alone, *KEAP1^KO^* cells developed G12Ci resistance more rapidly than NT controls, and *KEAP1*/*STK11 DKO* cells were intrinsically resistant to 10 nM adagrasib (**Fig. 6F**) or 30 nM sotorasib (**Fig. 6G**) confirming that the H358 KO panel recapitulates the reduced responsiveness seen in patients whose tumors harbor *KEAP1* ± *STK11* co-mutations. Intriguingly, SOS1i completely inhibited the development of G12Ci resistance in *KEAP1^KO^* cells and showed a significant enhancement of the treatment window in *KEAP1*/*STK11 DKO* cells (**Fig. 6H**), indicating that SOS1i has the potential to delay the development of G12Ci acquired resistance in the setting of tumors with *KEAP1* ± *STK11* co-mutations. Taken together, these data illustrate that co-mutations may not limit the usefulness of G12Ci + SOS1i combination therapy.

## Discussion

Mutations in KRAS^G12C^ are responsible for nearly 13% of LUAD cases (2). For patients with *KRAS^G12C^*-mutated LUAD, KRASG12C inhibitors have enormous therapeutic potential, however, both intrinsic and acquired resistance limit their overall effectiveness (4,13–17,23,27–30).

Studies show that inhibition of proximal RTK signaling using SHP2 (3,6,18,19,80) or SOS1 (51,53) inhibitors enhances G12Ci binding to mutated KRAS^G12C^ and limits RTK-SOS-WT RAS signaling to overcome intrinsic G12Ci resistance, and, as a result, enhances G12Ci efficacy *in vitro* and *in vivo*. Here, we show that SOS1 signaling lies at the crossroad of intrinsic and acquired G12Ci resistance. SOS1i enhanced the potency of G12Ci and inhibited rebound RTK-SOS-WT RAS signaling to limit intrinsic resistance in a SOS2-dependent manner. SOS1i further re-sensitized drug tolerant persister cells to G12Ci, thereby prolonging the window of G12Ci effectiveness and limiting the frequency of acquired G12Ci resistance.

While resistance to targeted therapies is generally framed as being either primary / intrinsic or secondary / acquired, emerging evidence suggests that for RTK/RAS pathway targeted therapies, resistance should be evaluated on a continuum based on the survival and evolution of DTPs (**Fig. 7**) (81). Understanding this continuum is key to understanding G12Ci responses in patients with co-mutations in *KEAP1* and *STK11* (16). Unlike primary or secondary resistance due to genetic mutations that directly circumvent G12Ci, DTPs use non-genetic chromatin remodeling that up-regulate signaling through multiple RTKs (35), enhance ability of cells to detoxify therapy-induced redox stress (36–40), and allow cells to enter a therapy-resistant near-quiescent state (33–35,41). Within the DTP population, TICs are capable of self-renewal and are hypothesized to represent the sanctuary population responsible for tumor recurrence after treatment failure (42,44). KEAP1 is a negative regulator of the NRF2 transcription factor and *KEAP1* loss-of-function results in enhanced NRF2 metabolic alterations leading to inability to regulate responses to oxidative damage (26,79). Patients with *KRAS^G12C^*-mutated LUAD harboring *KEAP1* co-mutations have inferior clinical outcomes and more often reach progression free survival of less than 3 months (16). While this could be interpreted as indicative of intrinsic resistance, an alternative hypothesis is that *KEAP1* LOF enhances DTP/TIC survival thereby limiting the overall effectiveness of G12Ci. Indeed, we observed that isogenic *KEAP1^KO^* cells showed an increased frequency of TICs (**Fig. 4**) and accelerated the development of acquired G12Ci resistance *in situ* (**Fig. 6**). These data suggest that patients whose tumors harbor *KEAP1* loss-of-function mutations may show resistance due to enrichment of the DTP population.

**Figure 7.**
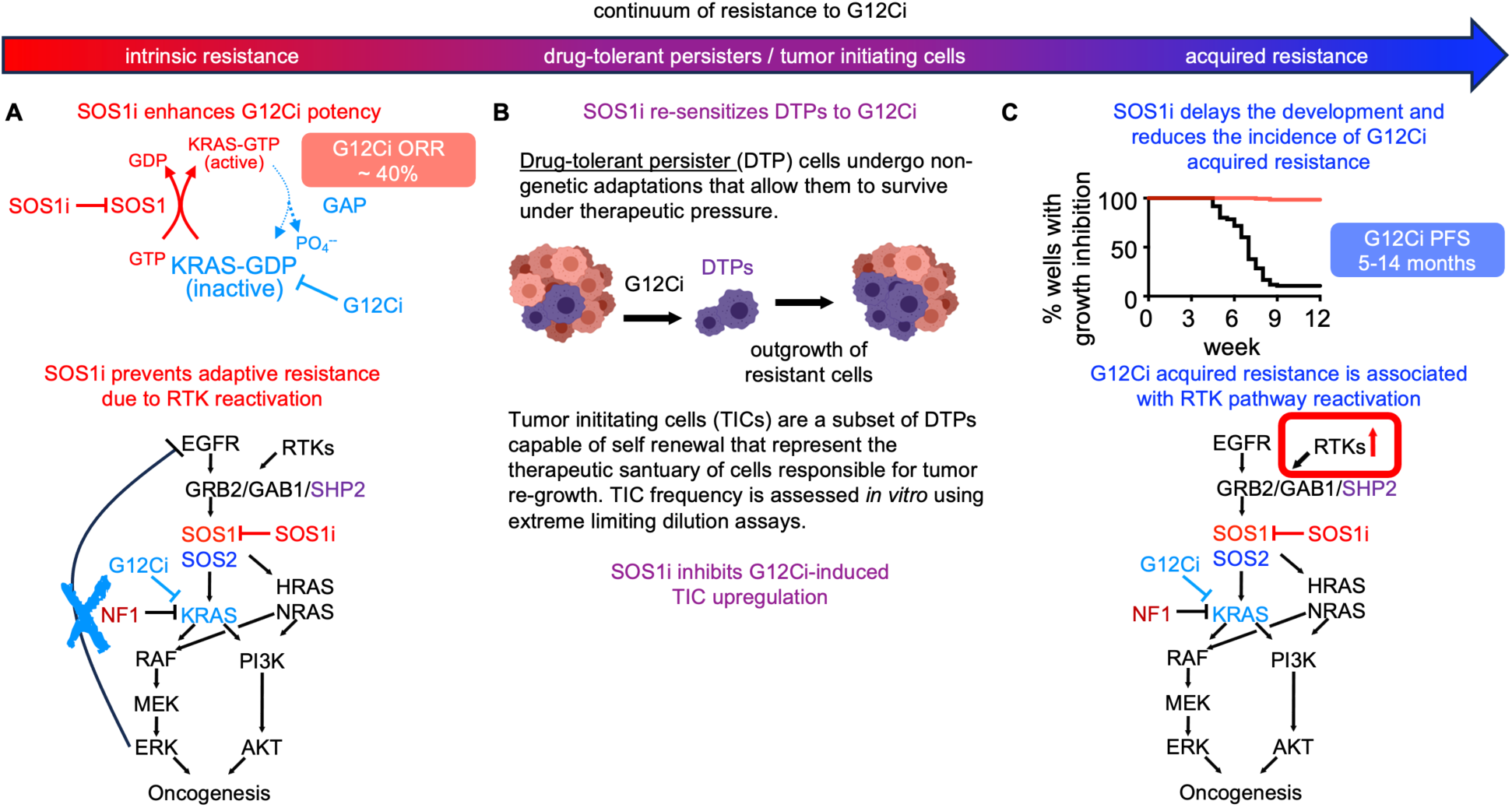
SOS1i targets the continuum of G12Ci resistant states. **A.** Intrinsic G12Ci resistance is driven by adaptive reactivation of RTK signaling due to a loss of ERK-dependent negative feedback. SOS1i targets rebound RTK signaling to limit adaptive G12Ci resistance. **B.** Cancer cells undergo non-genetic adaptation to G12Ci to both alter the redox environment and enhance alternative RTK signaling, both of which allow these ‘drug tolerant persister’ (DTP) cells to survive under therapeutic pressure. Within the DTP population, a subset of ‘tumor initiating cells’ (TICs) are capable of self-renewal and are thought to be the pharmacologic sanctuary driving that ultimately develop acquired resistance. SOS1i re-sensitizes DTPs to G12Ci and reduces TIC frequeny in G12Ci treated cultures. **C.** Acquired G12Ci resistance is often driven by RTK/RAS pathway reactivation by both genetic and non-genetic mechanisms. SOS1i both delayed the development of and reduced the frequency with which cultures acquired G12Ci resistance.

Unbiased assessment of signaling pathways modulated by combined SOS1i:G12Ci revealed that SOS1i inhibited both RTK/MAPK and hypoxia/HIF1α pathways in G12Ci treated cells (**Fig. 1**). Intriguingly, the combination of SOS1i-dependent inhibition of RTK signaling and hypoxia-associated pathways may underly SOS1i-dependent inhibition of both DTP/TIC survival (**Figs. 3 and 5**) and acquired G1Ci resistance (**Fig. 5**). Following G12Ci treatment, loss of ERK-dependent negative feedback on multiple RTKs drives adaptive G12Ci resistance and chromatin remodeling of DTPs activates multiple RTKs; HIF1α promotes transcription of genes responsible for TIC activity (71,72) and hypoxia signatures are associated with cancer stemness (42,63,82,83), DTP survival (62–64), and poor survival for patients with LUAD (73,74). SOS1i both re-sensitized DTPs to G12Ci (**Fig. 3**) and inhibited G12Ci-induced DTP survival (**Fig. 4**), likely due to the combined effects of SOS1i inhibiting RTK signaling and hypoxia-associated pathways in DTPs. Previous work shows that *Sos1*^-/-^ increases mitochondrial oxidative stress in MEFs due to mitochondrial dysfunction (84,85) so that SOS1 is an important regulator of redox signaling. Increased oxidative stress can lead to ROS accumulation to drive both senescence and multiple forms of cell death including ferroptosis (86). This oxidative stress is counteracted by KEAP1-dependent degradation of NRF2; tumors with *KEAP1* inactivating mutations show increased NRF2 activity and preventing the initiation of ferroptosis (87). NRF2 activity is further enhanced in cells harboring *STK11*/*KEAP1* co-mutations compared to KEAP1 mutations alone, suggesting that patients whose tumors harbor *KEAP1* ± *STK11* inactivating mutations are particularly insensitive to oxidative stress caused by targeted therapeutics. We found that SOS1i reduced DTPs and inhibited G12Ci resistance in *KEAP1^KO^* and *KEAP1/STK11 DKO* cells (**Fig. 6**). SOS1i may similarly tip the balance of oxidative stress in KEAP1i LOF and/or G12Ci-treated cells to inhibit DTPs survival, thereby enhancing the effectiveness of and blocking acquired resistance to G12Ci for patients whose tumors harbor *KEAP1* LOF mutations.

G12Ci effectiveness is limited by intrinsic and acquired resistance, hence, combination approaches are imperative to enhance clinical outcomes for patients with *KRAS^G12C^*-mutated tumors. Our study provides a framework for modeling the evolution of G12Ci resistance and assessing the impact of therapeutic combinations during this evolution. Our data suggests that SOS1i can limit resistance at each step of G12Ci resistance (**Fig. 7**). SOS1i enhanced G12Ci potency and inhibited adaptive resistance to increase G12Ci killing of LUAD cells. SOS1i further re-sensitized DTPs to G12Ci to delay and inhibit the development of acquired G12Ci resistance *in situ*. Our finding that SOS1i can target each stage of G12Ci resistance, even in the presence of KEAP1/STK11 co-mutations that predict insensitivity to G12Ci alone, reveals the therapeutic versatility of SOS1i + G12Ci combination therapy.

## Methods

### Cell Culture

Lung cancer cell lines were purchased from ATCC. Once cell lines were received, they were expanded and frozen at passage 3 and 4; cells were passaged once they became approximately 80% confluent. Cells were maintained in culture for 2-3 months before a new vial was thawed since prolonged passaging can alter TIC frequency (88). The panel of H358 NT, *STK11 ^KO^*, *KEAP1^KO^*, and *STK11*/*KEAP1 DKO* cells were a generous gift from Charles Rudin (Memorial Sloan Kettering Cancer Center) (79). Cell lines were cultured at 37°C and 5% CO2. All cells were passaged in RPMI supplemented with 10% FBS and 1% penicillin/streptomycin. For 3D signaling experiments, cells were seeded in a 24-well micropatterned Aggrewell 400 low-attachment plates (StemCell) at 9 x 10^5^ cells/well in 2 mL of medium. After 24 hrs post-plating, 1 mL of media was removed to be replaced with either new media or 2x inhibitor. Cells were treated for 0-72 hours in the same manner, replacing 1 mL of media every day with either new media or 2x inhibitor, or fresh 1x inhibitor for those that previously were drugged.

### Generation of *SOS2^KO^* cell lines

For *SOS2^KO^* studies, H358, H1373, H1792, and H2030 cells were infected with lentiviruses based on pLentiCRISPRv2 with either a non-targeting sgRNA (NT) or a sgRNA targeting *SOS2* (55). After 48 hrs post-infection, 4 μg/mL Puromycin (Invitrogen) was used to select cells. These cells were then analyzed for SOS1 or SOS2 expression after 7-10 days post-selection and only those cultures showing >80% deletion were used for the studies.

### Bliss Independence Analysis and MuSyC for Synergy

Cells were seeded at 750 cells per well in 100 μL in the inner-60 wells of 96-well ultra-low attachment round bottomed plates (FaCellitate BIOFLOAT # F202003) and allowed to coalesce as spheroids for 24-48h prior to drug treatment. For all studies, peripheral wells (rows A and H, columns 1 and 12) were filled with 200 μL of PBS to buffer inner wells and prevent evaporative effects on cells. Triplicate wells were then treated with increasing doses of adagrasib or BI-3406 alone or a combination of adagrasib + BI-3406 in a 9 × 9 matrix of drug combinations on a semi-log scale for 96h. and assessed for cell viability using CellTiter-Glo 2.0 (30 μL/well).

Luminescence was assessed using a Bio-Tek Cytation five multi-mode plate reader. Data were normalized to the maximum luminescence reading of untreated cells, and individual drug EC_50_ values were calculated using Prism9 by non-linear regression using log(inhibitor) vs. response. For all drug-treatment studies, the untreated sample for each cell line was set to 100%. This would mask any differences in 3D cell proliferation seen between cell lines. Excess over Bliss was calculated as the Actual Effect – Expected Effect as outlined in (89). The SUM EOB is calculated by taking the sum of excess over bliss values across the 9 x 9 treatment matrix. EOB values > 0 indicate increasing synergy. To deconvolute synergistic synergy versus potency, data were analyzed by Multi-dimensional Synergy of Combinations (MuSyC) Analysis (90,91) using an online tool (https://musyc.lolab.xyz) and are presented as mean +/- 95% confidence interval.

### Cell lysis and Western blotting

To prepare for cell lysis, cells were washed twice with cold PBS before RIPA buffer (1% NP-40, 0.1% SDS, 0.1% Na-deoxycholate, 10% glycerol, 0.137 M NaCl, 20 mM Tris pH [8.0], protease (Biotool #B14002) and phosphatase (Biotool #B15002) inhibitor cocktails) was added to cells for 20 minutes at 4°C and spun at 10,000 RPM for 10 min. Supernatant was collected and protein concentration was assessed via protein assay. Normalized lysates were then boiled in SDS sample buffer containing 100 mM DTT for 10 min prior to Western blotting. Proteins were resolved by sodium dodecyl sulfate-polyacrylamide (Criterion TGX precast) gel electrophoresis and transferred to nitrocellulose membranes. Western blots were developed by multiplex Western blotting using anti-SOS1 (Santa Cruz sc-256; 1:500), anti-SOS2 (Santa Cruz sc-258; 1:500), anti-β-actin (Sigma AC-15; 1:5,000 or Santa Cruz Biotechnology sc-47778, 1:2000 dilution), anti-pERK1/2 (Cell Signaling 4370; 1:1,000), anti-ERK1/2 (Cell Signaling 4696; 1:1000) primary antibodies. Anti-mouse and anti-rabbit secondary antibodies conjugated to IRDye680 or IRDye800 (LI-COR; 1:20,000) were used to probe primary antibodies. Western blot protein bands were detected and quantified using the Odyssey system (LI-COR).

### Transcriptome profiling

Gene expression (RNA-seq) analysis was performed as previously described (52). NT and *SOS2^KO^* H358 cells were seeded in 6-well micropatterned Aggrewell 400 low-attachment plates (StemCell) at 3.5 × 10^6^ cells/well in 4 mL of medium. Half of the media was removed and fresh media was added daily. 72h post-plating, 2 mL of media was removed to be replaced with 2 × SOS1i, G12Ci, or SOS1i + G12Ci and incubated for either 6h or 72h. Cells were collected, pelleted, resuspended in RLT plus buffer (Invitrogen), and homogenized using a QIAshredder column (Qiagen). RNA was isolated using an RNeasy spin column (Qiagen) according to the manufacturer’s instructions. After elution, RNA was quantified with the Quant-iT Ribogreen RNA Kit according to the manufacturer’s instructions (Invitrogen). All samples were normalized to 20 ng/μL, aliquoted and frozen at -80°C For each sample, a total RNA input of 200 ng was used for library preparation using the Illumina Stranded Total RNA Prep with Ribo-Zero Plus Kit (Illumina) and IDT for Illumina RNA UD Indexes Set A, Ligation (96 Indexes, 96 Samples). Sequencing libraries were quantified by real-time PCR using KAPA Library Quantification Kit for NGS (Roche) and assessed for size distribution on a Fragment Analyzer (Agilent) to confirm absence of adapter dimers and unligated library molecules. Sequencing libraries were pooled at <40-plex and sequenced on a NovaSeq 6000 (Illumina) using a 200 cycle SBS kit (Illumina) with paired-end reads at 101 bp length targeting >60 million reads per sample.

After sequencing, the Nextflow nf-core/rnaseq (version 3.4) workflow was used to estimate a matrix of sample gene expression values (92–94) using the reference human genome (GRCh37; Ensembl release 75) obtained from Illumina’s iGenomes resource (https://support.illumina.com/sequencing/sequencing_software/igenome.html). Briefly, paired-end DNA sequence FastQ files were merged by sample (two/sample) and underwent read quality control (FastQC v0.11.9) prior to adapter and quality trimming (Trim Galore! v0.6.7), then input into Salmon’s (v1.5.2) (95) mapping-mode for gene expression matrix quantification.

Further sequencing quality control was performed via subsequent trimmed read alignment using STAR (v2.6.1d) (96), SAMtools (v1.13) (97), and Picard (v2.25.7, http://broadinstitute.github.io/picard/), followed by RSeQC (v3.0.1) (98), Qualimap (v2.2.2-dev) (99), and Preseq (v3.1.1, https://smithlabresearch.org/software/preseq/). After confirming the absence of sequencing or sequencing library anomalies, we proceeded to differential expression analysis.

Gene set enrichment analysis was performed using fgsea (100) R/Bioconductor (v1.14) package and hallmark gene sets from the molecular signatures database (MSigDB v7.5.1) (60). The resulting nominal p values were adjusted using Benjamini-Hochberg multiple testing correction method and gene sets with adjusted p value < 0.01 were considered as significant.

Estimation of individual transcription factor activities based on ‘footprint based’ signatures for individual transcription factors were performed using the Discriminant Regulon Expression Analysis (DoRothEA) R/Bioconductor package (101,102) and Visualization Pipeline for RNA-seq (VIPER) analysis (103). The Pathway RespOnsive GENes (PROGENy) algorithm was used to infer activation of 14 key cancer pathways (61). MAPK Pathway Activity (MPAS) scores were calculated based on expression of the 10-MAPK gene signature (59). Heatmaps of GSEA analysis were generated using ComplexHeatmap (v2.4.3) (104) or Prism 9. All other figures from data analyses were visualized using ggplot2 (v3.3.2) (105) R package.

### Flow cytometry

Cells were plated in 6 cm tissue-culture treated plates and allowed to adhere for 24 hrs prior to adding G12Ci at the indicated doses for 72 hrs. After 72 hrs of treatment, cells were harvested by trypsinization, spun down, resuspended in Aldefluor Assay Buffer (StemCell) at 1*10^6^ cells/mL, and stained for ALDH activity using the Aldefluor Assay Kit per manufacturer’s instructions. Two tubes of untreated cells was collected so one could be treated with DEAB, an ALDH inhibitor, to use as a negative gating control during analysis. Data was analyzed using FloJo with and are presented as the percentage of cells showing ALDH activity over DEAB controls.

### Extreme Limiting Dilution Assays

For an *in situ* assay, cells are seeded in a 96-well ultra-low attachment flat bottomed plates (Corning Corstar #3474). Cells are plated at decreasing cell concentrations at half-log interavals, beginning with 1000 cells/well – 1 cell/well, with 12 wells/cell concentration except 10 cells/well, where 24 wells are seeded. To assess the effect of G12C inhibition on TIC frequency, cells were left untreated or were pre-treated with the indicated doses of G12Ci for 72-h, after which drugged media was removed and replaced with fresh media for 48-72 hrs prior to plating. To assess the effect of SOS1 inhibition on TICs, cells were seeded ± 300 nM BI-3406. Cells were then allowed to grow for 7-10 days before being assessed for spheroid growth (>100 mM was scored as positive). The TIC frequency and significance between the different groups was calculated by ELDA website https://bioinf.wehi.edu.au/software/elda/ (106).

### Resistance Assays

Cells were plated in 96-well tissue-culture treated plates at low-density (250 cells/well). Each plate was treated with a single dose of a G12C inhibitor ± BI-3406. Wells were assessed weekly for outgrowth and those that were >50% confluent were marked as resistant to that given dose. Plates that did not reach full confluency were fed weekly until an indicated endpoint was reached. Data are plotted as a Kaplan-Meyer survival curve; significance was assessed by comparing Kaplan-Meyer curves using Prism 9.

## Data availability

The data generated in this study are available upon request from the corresponding author.

## Acknowledgments

We thank the USUHS Bioinformatics Core. We thank Charles Rudin (Memorial Sloan Kettering Cancer Center) for *STK11* and *KEAP1* KO cells. **Funding**: This work was supported by funding from the NIH (R01 CA255232 and R21 CA267515 to R.L.K.) and a CRADA from Boehringer Ingelheim (to R.L.K). The funders had no role in the study design, data collection and interpretation, or the decision to submit the work for publication. The opinions and assertions expressed herein are those of the authors and are not to be construed as reflecting the views of Uniformed Services University of the Health Sciences or the United States Department of Defense. **Competing interests:** The Kortum laboratory receives funding from Boehringer Ingelheim to study SOS1 as a therapeutic target in *RAS*-mutated cancers.

---

**Figure S6 (related to Fig. 5).**
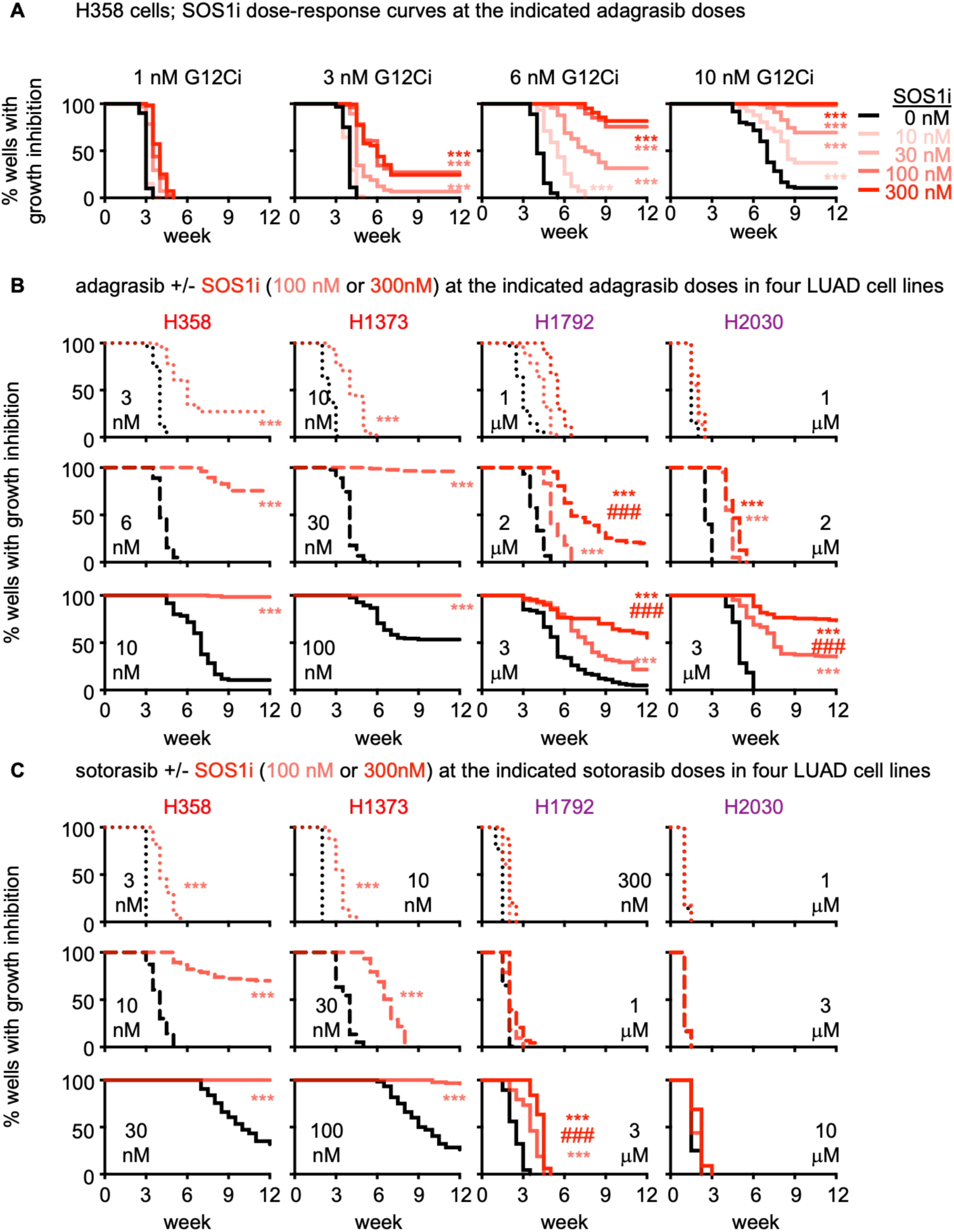
SOS1 inhibition limits the development of acquired G12Ci resistance. Multi-well resistance assays were performed as outlined in the Materials and Methods. **A.** G12Ci resistance in H358 cells treated with the indicated dose of adagrasib alone (black) or increasing doses of SOS1i (reds). **B-C**. G12Ci resistance to the indicated dose of adagrasib (B) or sotorasib (C) in H358, H1373, H1792, or H2030 cells treated with a low (dotted), intermediate (dashed), or high (solid) dose of the G12Ci adagrasib alone (black) or G12Ci + 100 nM (light red) or 300 nM (dark red) SOS1i. G12CI doses were based on the highest three doses that allowed the development of G12Ci resistance in each parental cell line. Data are pooled from three independent experiments. *** p < 0.001 vs. G12Ci alone; ### p < 0.001 for cells treated with 100 vs 300 nM SOS1i.

## Notes

### Summary of Updates

Corrected author spelling.

