## Supplementary Figures for "SOS1 inhibition enhances the efficacy of and delays resistance to G12C inhibitors in lung adenocarcinoma"

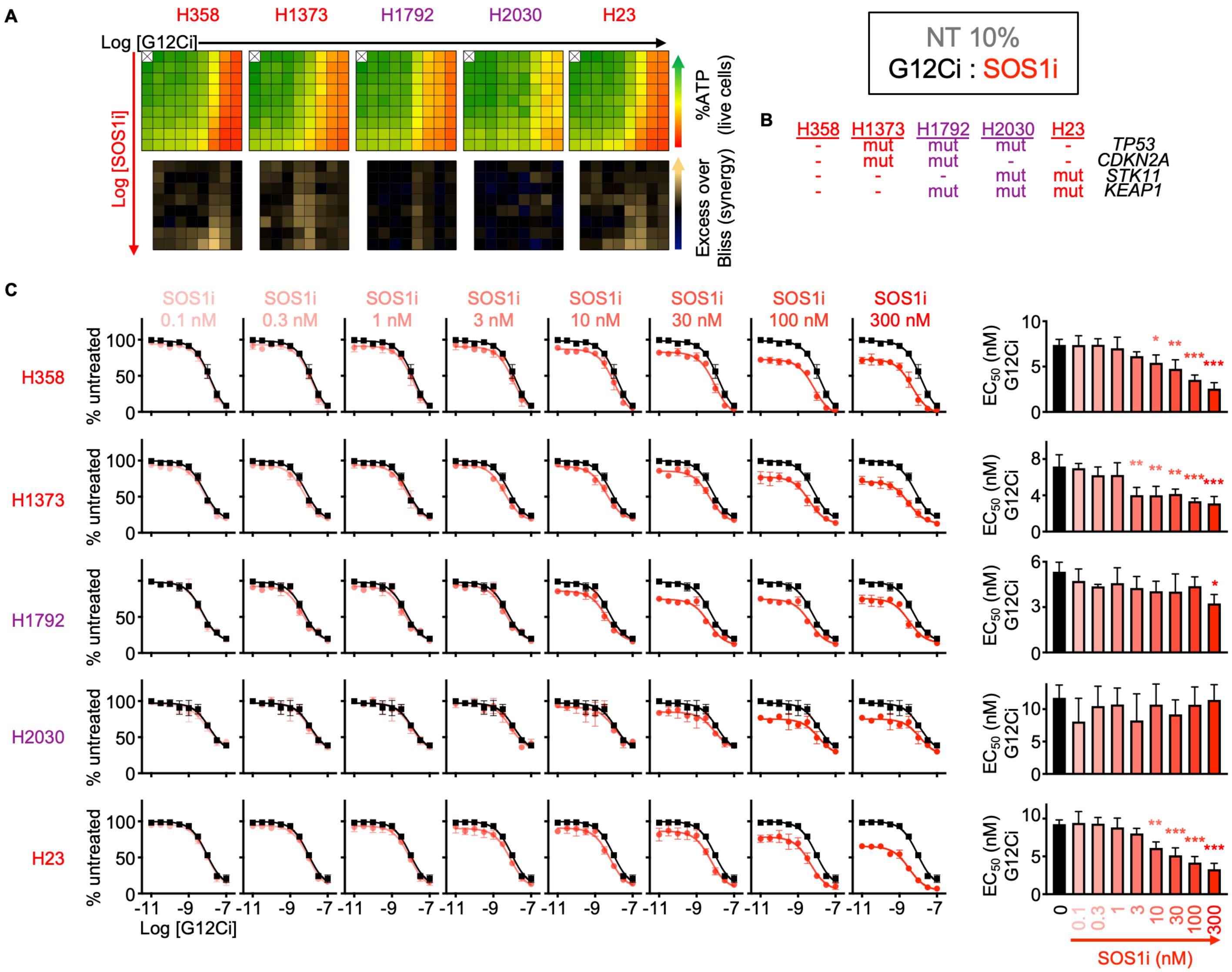


**Figure S1 (related to Fig. 1A-C)**. LUAD cells show varying SOS1i:G12Ci synergy in 10% serum.

**A**. Heat map of cell viability (top) and excess over Bliss (EOB, bottom) for the indicated *KRAS*$^{G12C}$-mutated LUAD cell lines treated with increasing (semi-log) doses of the G12Ci adagrasib ($10^{-10.5}$ – $10^{-7}$), the SOS1i BI-3406 ($10^{-10}$ – $10^{-6.5}$) or the combination of G12Ci + SOS1i under 3D spheroid culture conditions in 10% serum. Data are the mean from three independent experiments, each experiment had three technical replicates. Data are repeated from Fig. 1A.

**B**. *TP53*, *CDKN2A*, *STK11*, and *KEAP1* mutation status in the LUAD cell lines from A.

**C**. G12Ci single-dose response curve indicating % cell viability for *KRAS*-mutated LUAD cell lines from A in anchorage-independent (3D) conditions for 72 hours in 10% serum.

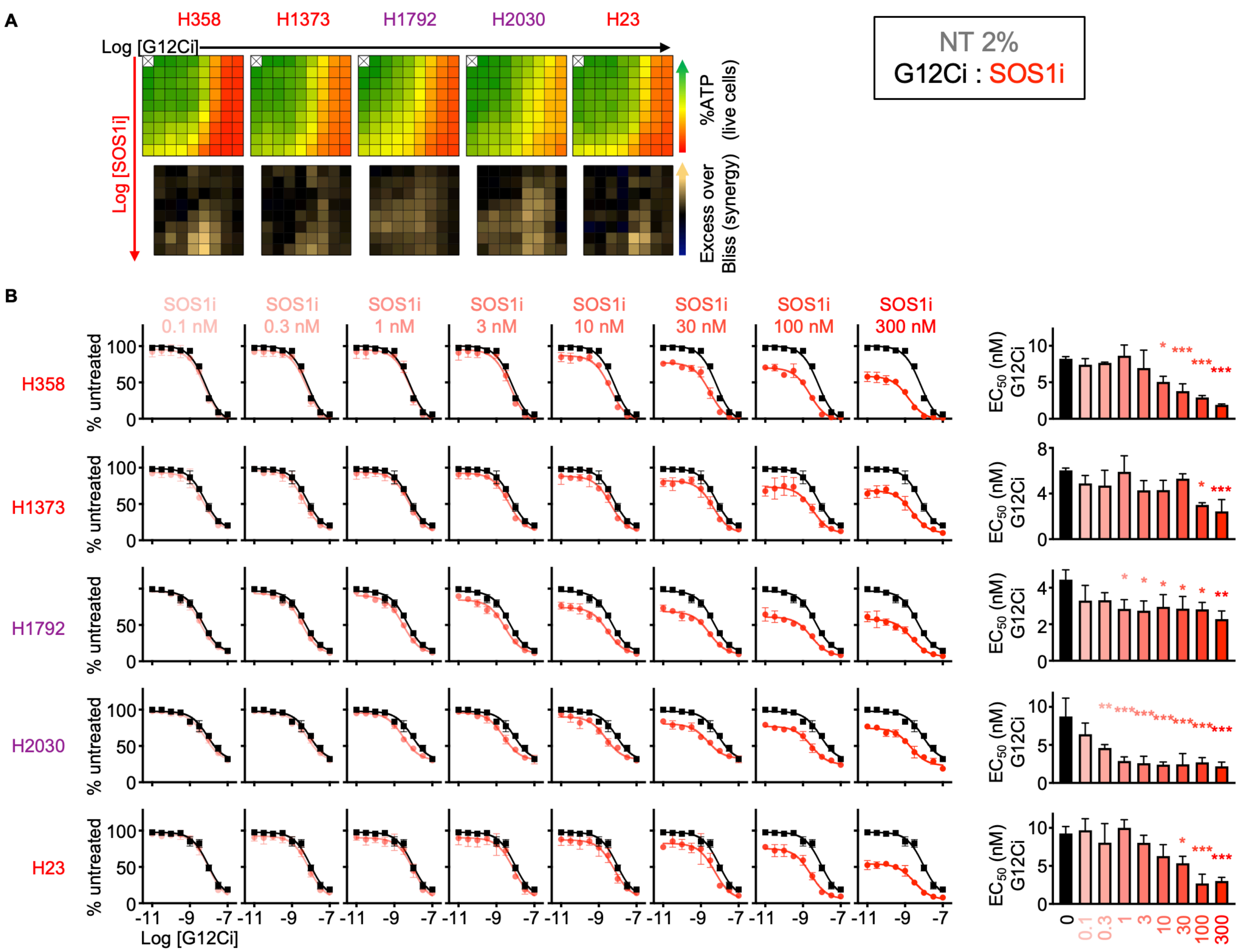


**Figure S2 (related to Fig. 1B-C)**. SOS1i:G12Ci synergy is enhanced by low serum culture. **A**. Heat map of cell viability (top) and excess over Bliss (EOB, bottom) for the indicated *KRAS*$^{G12C}$-mutated LUAD cell lines treated with increasing (semi-log) doses of the G12Ci adagrasib ($10^{-10.5}$ – $10^{-7}$), the SOS1i BI-3406 ($10^{-10}$ – $10^{-6.5}$) or the combination of G12Ci + SOS1i under 3D spheroid culture conditions in 2% serum. Data are the mean from three independent experiments, each experiment had three technical replicates.
**B**. G12Ci single-dose response curve indicating % cell viability for *KRAS*-mutated LUAD cell lines from A in anchorage-independent (3D) conditions for 72 hours in 2% serum.

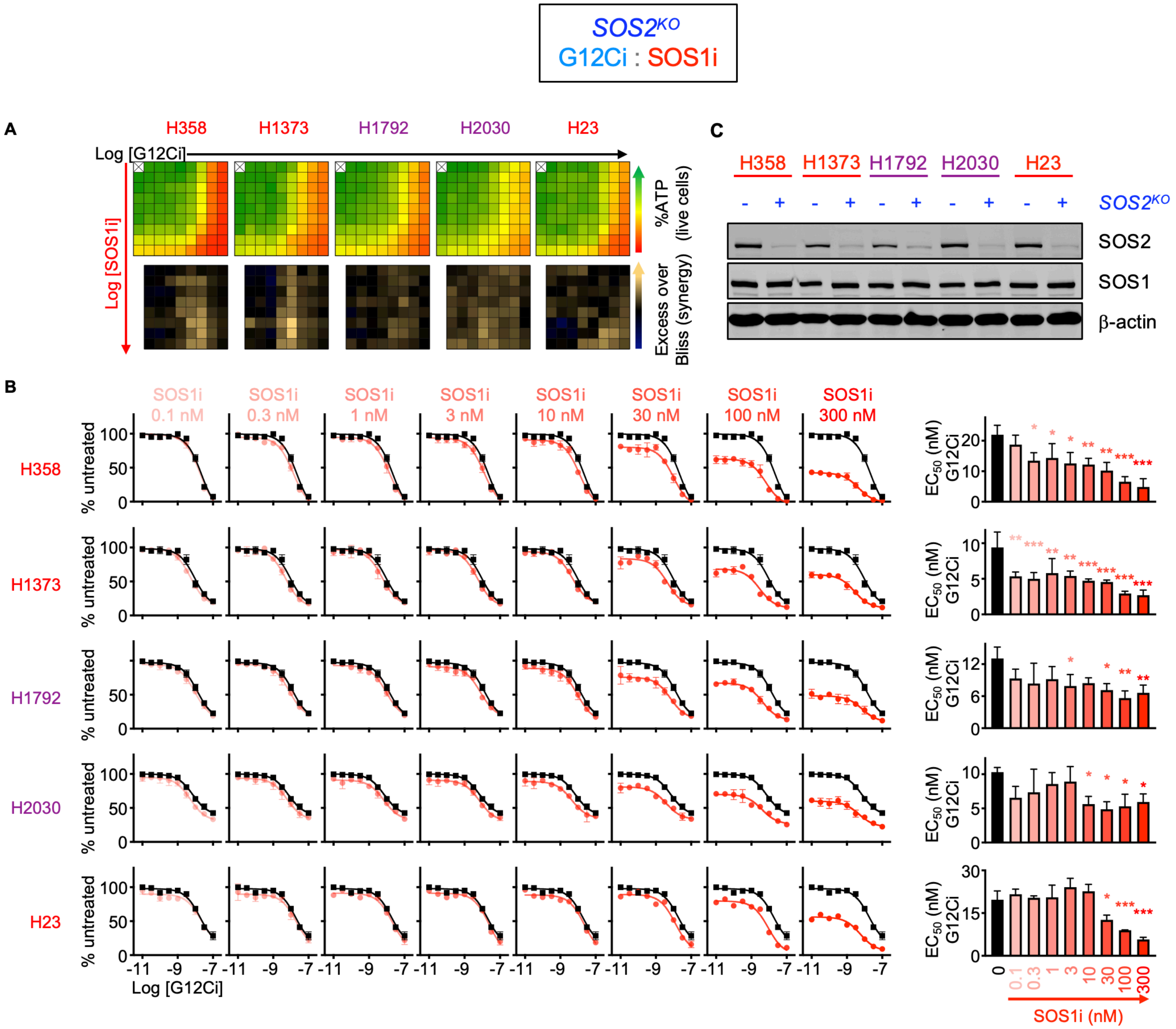


**Figure S3 (related to Fig. 1B-C)**. *SOS2*$^{KO}$ restores SOS1i:G12Ci synergy in 10% serum.

**A**. Heat map of cell viability (top) and excess over Bliss (EOB, bottom) for the indicated *SOS2*$^{KO}$ *KRAS*$^{G12C}$-mutated LUAD cell lines treated with increasing (semi-log) doses of the G12Ci adagrasib ($10^{-10.5}$ – $10^{-7}$), the SOS1i BI-3406 ($10^{-10}$ – $10^{-6.5}$) or the combination of G12Ci + SOS1i under 3D spheroid culture conditions in 10% serum. Data are the mean from three independent experiments, each experiment had three technical replicates.

**B**. G12Ci single-dose response curve indicating % cell viability for *KRAS*-mutated LUAD cell lines from A in anchorage-independent (3D) conditions for 72 hours in 2% serum.

**C**. Western blots for SOS2, SOS1, and b-actin showing *SOS2*$^{KO}$ (>80%) in pooled cells for the indicated LUAD cell line.

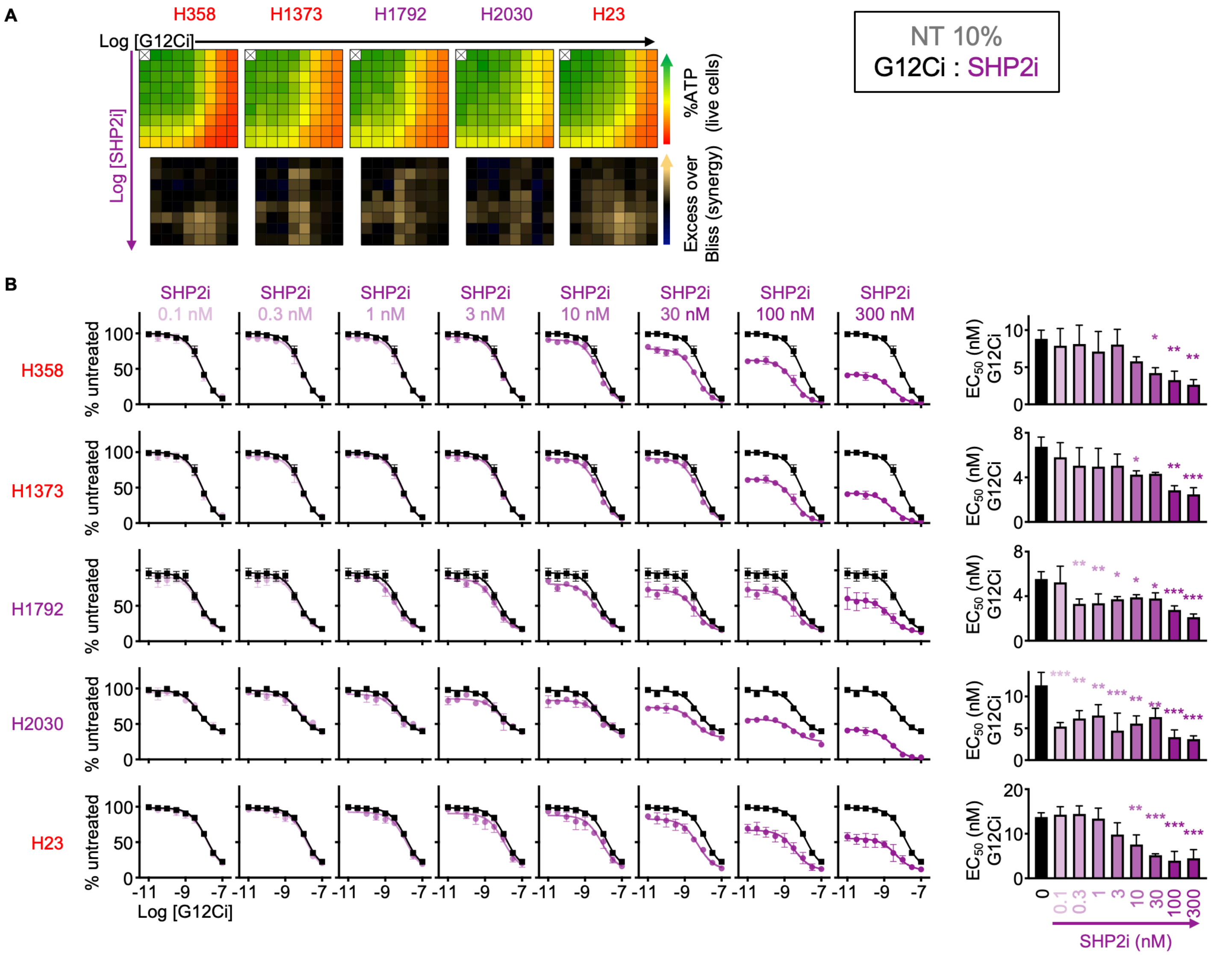


**Figure S4 (related to Fig. 1B-C)**. SHP2i:G12Ci synergy in LUAD cells.

**A**. Heat map of cell viability (top) and excess over Bliss (EOB, bottom) for the indicated *KRAS*$^{G12C}$-mutated LUAD cell lines treated with increasing (semi-log) doses of the G12Ci adagrasib ($10^{-10.5}$ – $10^{-7}$), the SHP2i RMC-4550 ($10^{-10}$ – $10^{-6.5}$) or the combination of G12Ci + SHP2i under 3D spheroid culture conditions in 10% serum. Data are the mean from three independent experiments, each experiment had three technical replicates.

**B**. G12Ci single-dose response curve indicating % cell viability for *KRAS*-mutated LUAD cell lines from A in anchorage-independent (3D) conditions for 72 hours in 2% serum.

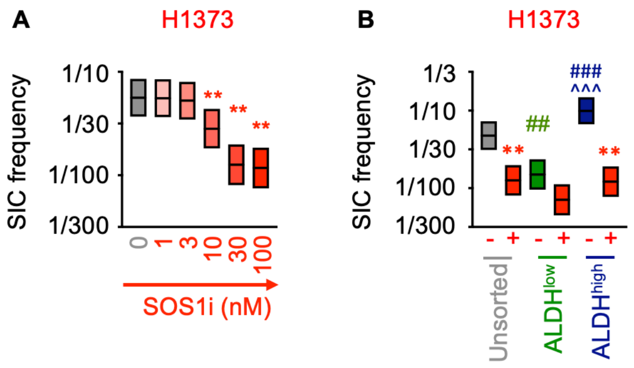


**Figure S5 (related to Fig. 4).** SOS1i ± *SOS2*$^{KO}$ prevents G12Ci-induced TIC outgrowth.

**A**. TIC frequency from *in situ* ELDAs of H1373 cells treated with the indicated SOS1i doses. * $\chi^2$ < 0.05, ** $\chi^2$ < 0.01 vs. NT untreated.

**B**. TIC frequency from *in situ* ELDAs in unsorted (grey), ALDH$^{low}$ (green), and ALDH$^{high}$ (dark blue) H1373 cells left untreated or treated with 100 nM BI-3406 (SOS1i). **$\chi^2$ < 0.01 vs untreated; ## $\chi^2$ < 0.01, ### $\chi^2$ < 0.001 vs. unsorted cells; ^^^ $\chi^2$ < 0.001 vs. ALDH$^{low}$ cells.

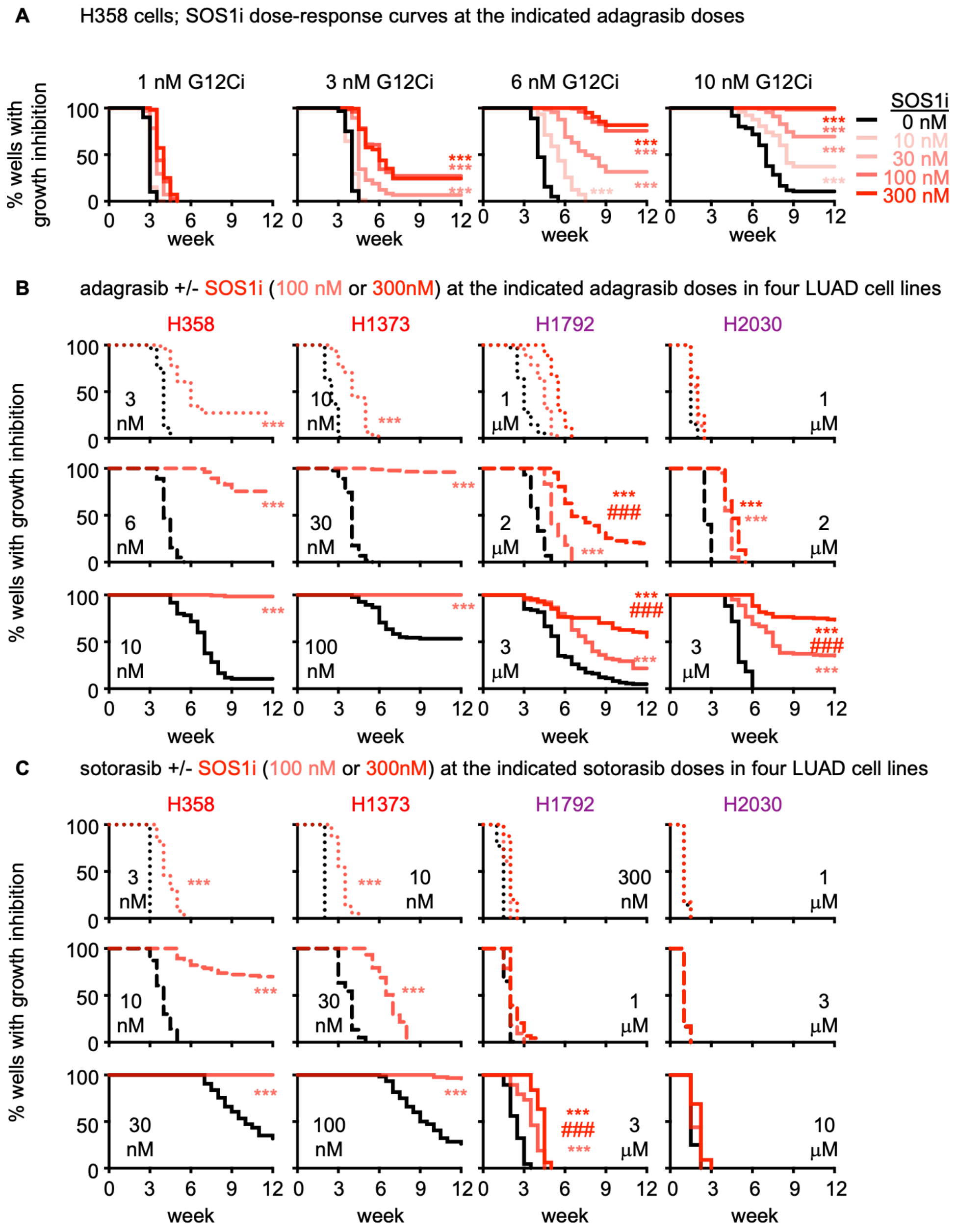


**Figure S6 (related to Fig. 5)**. SOS1 inhibition limits the development of acquired G12Ci resistance.

Multi-well resistance assays were performed as outlined in the Materials and Methods.

**A**. G12Ci resistance in H358 cells treated with the indicated dose of adagrasib alone (black) or increasing doses of SOS1i (reds).

**B-C**. G12Ci resistance to the indicated dose of adagrasib (B) or sotorasib (C) in H358, H1373, H1792, or H2030 cells treated with a low (dotted), intermediate (dashed), or high (solid) dose of the G12Ci adagrasib alone (black) or G12Ci + 100 nM (light red) or 300 nM (dark red) SOS1i. G12CI doses were based on the highest three doses that allowed the development of G12Ci resistance in each parental cell line. Data are pooled from three independent experiments. *** $p < 0.001$ vs. G12Ci alone; ### $p < 0.001$ for cells treated with 100 vs 300 nM SOS1i.
